# 3D Spatial Interactomics Maps the Dynamics of NF-κB Multiprotein Signalosomes in Single Cells

**DOI:** 10.64898/2026.09.14.751198

**Authors:** Nicholas Zhang, Collin Leese-Thompson, Hazel Ozuna, Beverly Peng, Sriya Sirigireddy, Dhruv Nambiar, Lakshana Ramanan, Rabindra Tirouvanziam, Benjamin Kopp, Ahmet F. Coskun

**Affiliations:** Wallace H. Coulter Department of Biomedical Engineering, Georgia Institute of Technology and Emory University, Atlanta, GA, USA; Interdisciplinary Bioengineering Graduate Program, Georgia Institute of Technology, Atlanta, GA, USA; Parker H. Petit Institute for Bioengineering and Bioscience, Georgia Institute of Technology, Atlanta, GA, USA; Department of Pediatrics, Emory University, Atlanta, GA, USA; Center for CF & Airways Disease Research, Children’s Healthcare of Atlanta, Atlanta, GA, USA; Center for Perinatal Research, The Abigail Wexner Research Institute at Nationwide Children’s Hospital, Columbus, OH, USA; Department of Pediatrics, The Ohio State University, Columbus, OH, USA

## Abstract

NFκB signaling drives inflammatory responses by rapidly assembling membrane-proximal multiprotein supercomplexes, yet how these assemblies are organized in space and time within the 3D interior of a single cell has remained uncharacterized. We addressed this by profiling endogenous NFκB protein-protein interactions with an intelligent sequential proximity ligation assay (iseqPLA), read out by spinning disk confocal microscopy and 3D reconstruction. Each detected protein-protein proximity event is represented by a rolling-circle amplification product, and we treat clusters of co-localized puncta as a measure of supercomplex spatial organization. Across NIH-3T3 mouse fibroblasts, cystic fibrosis (CF) patient-derived macrophage co-cultures with IMR-90 human fibroblasts, and an independent set of healthy-and CF-donor monocyte-fibroblast co-cultures profiled by 3D iseqPLA, we tracked supercomplex dissociation, p65 nuclear translocation, and negative-feedback engagement across cytokine time courses. Three findings emerge: 3D volumetric quantification reduces the variance in nuclear-to-cytoplasmic ratio measurements relative to 2D projections, the choice of extracellular matrix coating shapes the fraction of NFκB-responsive cells, and CF airway-conditioned macrophages amplify paracrine NFκB signaling in neighboring fibroblasts in a CF model. A single-cell Generative Pretrained Transformer (scGPT) foundation model, fine-tuned on curated transcriptomic datasets, further places our NFκB gene panel within an inflammation-relevant feature space. Together, these results establish a 3D spatial interactomics workflow for dissecting supercomplex dynamics in health and disease.

## Introduction

The nuclear factor kappa-light-chain-enhancer of activated B cells (NFκB) pathway is a central controller of inflammation, innate and adaptive immunity, cell survival, and proliferation ^[1,2]^. Since Sen and Baltimore first described it in 1986 ^[3]^, the family has been tied to chronic inflammatory diseases that include rheumatoid arthritis, inflammatory bowel disease, asthma, and cystic fibrosis (CF) ^[2,4,5]^. Proinflammatory cytokines, among them tumor necrosis factor alpha (TNFα) and interleukin-1 beta (IL-1β), engage their receptors to trigger the canonical pathway, a cascade that converges on the IκB kinase (IKK) complex ^[1,6]^. Activated IKK phosphorylates IκBα and marks it for proteasomal degradation, freeing the p65/p50 heterodimer to enter the nucleus and drive transcription of proinflammatory genes ^[1,2,6,7]^. This output is held in check by negative feedback: NFκB induces A20 (TNFAIP3), a deubiquitinase that dampens upstream signaling ^[5,8]^.

Initiating an NFκB signal requires the rapid assembly of multiprotein supercomplexes at membrane-proximal hubs. Following established usage in mitochondrial and signalosome biology, we use “supercomplex” to mean a higher-order assembly of individually stable protein complexes ^[9]^. In our imaging readout, we use the term for putative higher-order assemblies inferred from rolling-circle-amplification proximity products: individual amplification products (∼0.5–1 μm across) can merge into confluent signal at high interaction density, so connected-component counts undercount proximity events in this saturated regime. During TNFα signaling, the TNFR1-associated Complex I brings together TRADD, TRAF2, TRAF5, RIP1, and cIAP1/2, building the polyubiquitin scaffolds that recruit TAK1 and IKK ^[6,10–12]^. TurboID proximity labeling has shown that p65 alone contacts more than 350 proteins, a reminder that native supercomplexes are far more diverse than any one method can capture ^[13]^. Downstream propagation in turn depends on these upstream supercomplexes coming apart, including the release of TRAF2 and TRAF5 from TRADD ^[10]^. Yet despite this well-characterized biochemistry, built largely on co-immunoprecipitation and structural studies, the spatiotemporal dynamics of NFκB protein-protein interactions (PPIs) have never been visualized at the single-cell level in their native 3D intracellular setting^[14,15].^

The proximity ligation assay (PLA) reads out endogenous PPIs at single-molecule resolution, pairing oligonucleotide-conjugated antibodies with rolling circle amplification (RCA) ^[16–18]^, and the Duolink implementation is now a standard way to visualize PPIs in fixed cells ^[19,20]^. Proximity sequencing variants (Prox-seq, Sprox-seq) join PLA to single-cell RNA sequencing for multiplexed detection ^[21–23]^, but these largely address surface-accessible targets and cannot resolve intracellular PPIs inside native subcellular compartments. To reach the cell interior, our group introduced intelligent sequential PLA (iseqPLA), which reads one PPI per cycle in a single fluorescent channel and then strips and restains antibodies to build up a multiplexed intracellular panel ^[24]^. Coupling iseqPLA to spinning disk confocal microscopy and 3D reconstruction sidesteps a limitation of conventional 2D PLA, which collapses volumetric information and can blur nuclear signal together with perinuclear cytoplasmic signal ^[24–27]^.

The extracellular matrix (ECM) is not passive scaffolding; it shapes intracellular signaling through integrin-mediated mechanotransduction ^[28,29]^. Collagen I, for instance, alters how fibroblasts respond to cytokines ^[29–33]^, yet how the coating substrate shapes NFκB PPI dynamics has gone largely unexamined. In a separate disease context, cystic fibrosis, caused by CFTR mutations, features dysregulated NFκB signaling and excessive proinflammatory cytokine production ^[5,34–36]^. Co-cultures built with CF patient-derived macrophages offer a tractable way to study paracrine NFκB signaling ^[37]^, and the Tirouvanziam laboratory’s organotypic airway transmigration model supplies the disease-conditioned macrophages such studies require ^[38,39]^.

Here we apply 3D iseqPLA to NFκB supercomplexes across three model systems: (1) NIH-3T3 mouse fibroblasts stimulated with TNFα and IL-1β, where we track supercomplex dissociation, p65 translocation, and A20-mediated feedback; (2) CF macrophage and monocyte co-cultures with IMR-90 human fibroblasts, where we probe paracrine NFκB amplification; and (3) a scGPT foundation model analysis ^[40]^, which tests whether our NFκB gene panel is enriched in inflammation-relevant transcriptional space. Across these systems, ECM coating shapes the fraction of responsive cells, CF macrophages amplify paracrine signaling, and 3D imaging yields lower measurement variance than 2D analysis.

## Results

### 3D iseqPLA profiles intracellular NFκB supercomplexes in situ

To resolve how intracellular NFκB supercomplexes are organized at single-cell resolution, we built a multiplexed imaging workflow that pairs iseqPLA with spinning disk confocal (SDC) microscopy and 3D volumetric reconstruction (Fig. 1a). Each imaging cycle captures a single PPI in one fluorescent channel (647 nm) using Duolink single-color proximity ligation; VectaPlex antibody stripping then clears that signal so the sample can be restained on the next cycle. Across up to four cycles we profiled the immunofluorescent reporter proteins H2B and p65 (Cycle 1), p105/p50 & p65 heterodimer formation (Cycle 2), A20 & IKKβ negative feedback (Cycle 3), and A20 & IKKγ (Cycle 4); a separate panel covered the upstream TRAF-5_TRADD and TRAF-5_TRAF-2 PPIs.

**Fig. 1.**
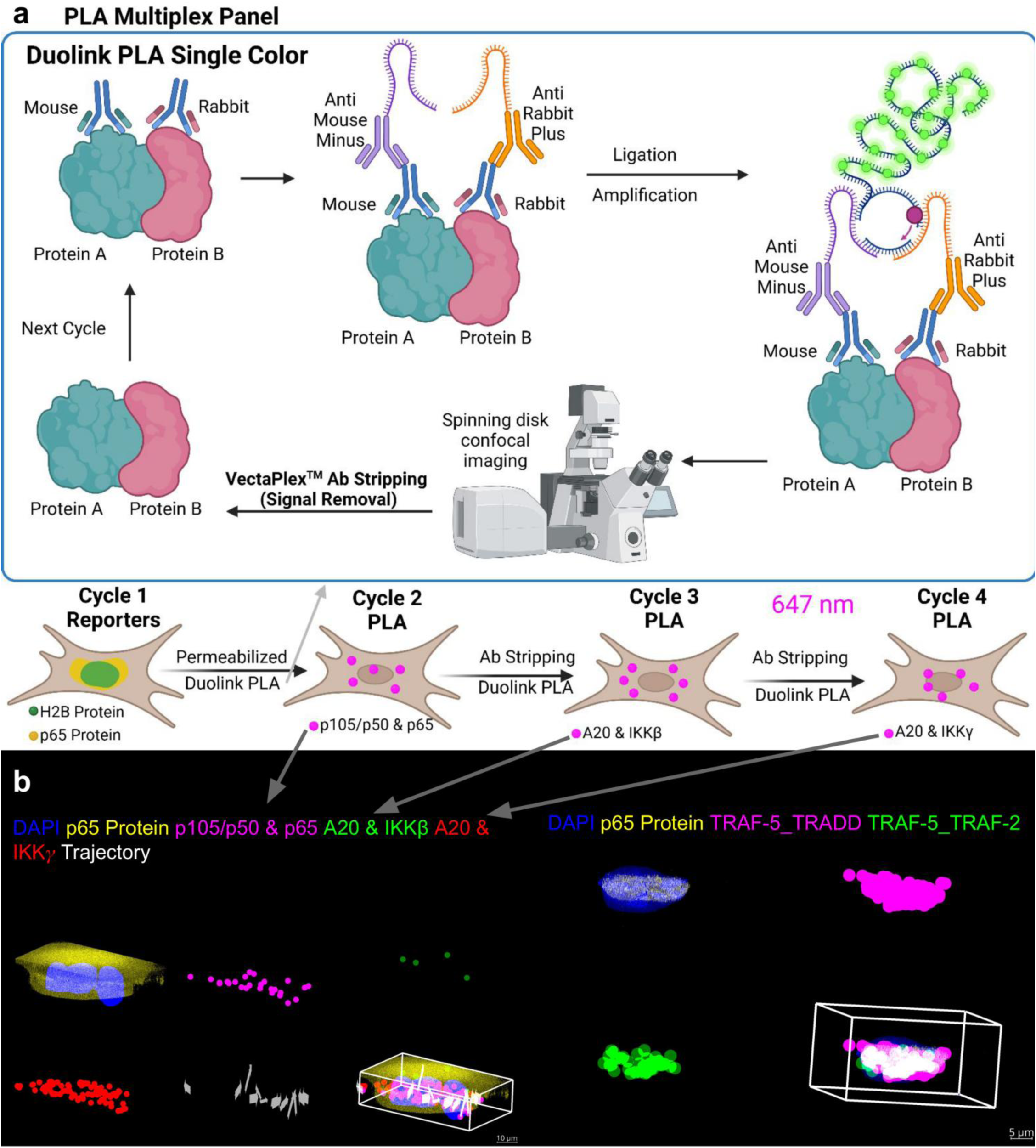
Multiplexed sequential PLA (iseqPLA) workflow for 3D in situ visualization of NFκB protein-protein interactions. **(a)** Schematic of the iseqPLA multiplex panel. Cycle 1 captures reporter proteins (H2B and p65 protein) via immunofluorescence in the A488/GFP and ds-RED/TRITC channels, respectively. Subsequent cycles (2–4) each detect a single PPI in the 647 nm channel using Duolink single-color PLA, targeting p105/p50 & p65 (Cycle 2), A20 & IKKβ (Cycle 3), and A20 & IKKγ (Cycle 4). Between cycles, antibody stripping with VectaPlex™ Signal Removal reagent removes prior PLA signal to enable sequential restaining. The Duolink PLA mechanism is illustrated: a pair of oligonucleotide-conjugated secondary antibodies (anti-mouse MINUS and anti-rabbit PLUS) bind to proximal primary antibody pairs targeting Protein A and Protein B; proximity-dependent ligation and rolling circle amplification generate a fluorescent DNA concatemer signal only when the two proteins are within ∼40 nm. Created with BioRender.com. **(b)** Representative 3D reconstructions from spinning disk confocal z-stacks showing multiplexed PLA signal in single cells. Left panels display the full iseqPLA panel including DAPI (nuclear stain, blue), p65 protein (yellow), p105/p50 & p65 PLA (magenta), A20 & IKKβ PLA (green), A20 & IKKγ PLA (red), and PPI trajectory analysis. Right panels show a separate NIH-3T3 fibroblast stained for DAPI (blue), p65 protein (yellow), TRAF-5_TRADD PLA (cyan), and TRAF-5_TRAF-2 PLA (magenta), with a 3D bounding box rendering. Scale bars: 15 μm (left), 5 μm (right).

We acquired every field as a full z-stack, 40 planes at 0.5 μm spacing, on a Cephla Squid SDC fitted with a Nikon 60× water objective, and reconstructed the signals in 3D with the PRISMS workflow ^[27]^. The spatial properties we report (area, volume, count) describe the fluorescent RCA products, which stand in as proxies for the underlying PPIs; the volumes therefore reflect clusters of RCA product rather than the true dimensions of a native supercomplex.

Representative 3D renderings of individual NIH-3T3 fibroblasts carrying the full iseqPLA panel show multiple NFκB PPI signals coexisting in one cell, each at a distinct subcellular location (Fig. 1b). In a separate NIH-3T3 fibroblast, the upstream panel separated TRAF-5_TRADD (cyan) from TRAF-5_TRAF-2 (magenta) as spatially distinct cytoplasmic clusters sitting adjacent to the p65 protein signal (yellow); 3D bounding-box renderings placed both firmly outside the nucleus across the whole cell volume (Fig. 1b, right). These supercomplex puncta populated the cytoplasmic z-volume and did not cluster in any single focal plane. The white overlay (“PPI trajectory”) is a per-punctum direction-vector field: from each upstream interaction punctum it points toward the local average position of its nearest downstream interaction puncta, as defined in Methods.

### 3D volumetric quantification of NFκB PPI dynamics in NIH-3T3 fibroblasts

To gauge how imaging dimensionality affects the readout, we computed nuclear-to-cytoplasmic (N/C) p65 ratios two ways, from 2D maximum intensity projections (MIP) and from 3D volumetric segmentation, after IL-1β stimulation of NIH-3T3 fibroblasts (Fig. 2b). This comparison drew on 2,780 cells imaged across all timepoints and conditions. The 2D route compressed the dynamic range, inflated cell-to-cell variance, and exaggerated N/C ratios at peak activation (maximum N/C ∼3 AU in 2D versus ∼2.5 AU in 3D), whereas 3D segmentation gave steadier trajectories with tighter variance. This happens because 2D projections stack nuclear and perinuclear cytoplasmic signal from many z-planes into one plane. Translocation kinetics look qualitatively similar either way, but 3D draws a cleaner line between nuclear and cytoplasmic signal. Because both metrics derive from the same cells and the same 4′,6-diamidino-2-phenylindole (DAPI)-based segmentation and differ only in projection dimensionality, the reduced variance reflects dimensionality rather than differences in sampling or segmentation.

**Fig. 2.**
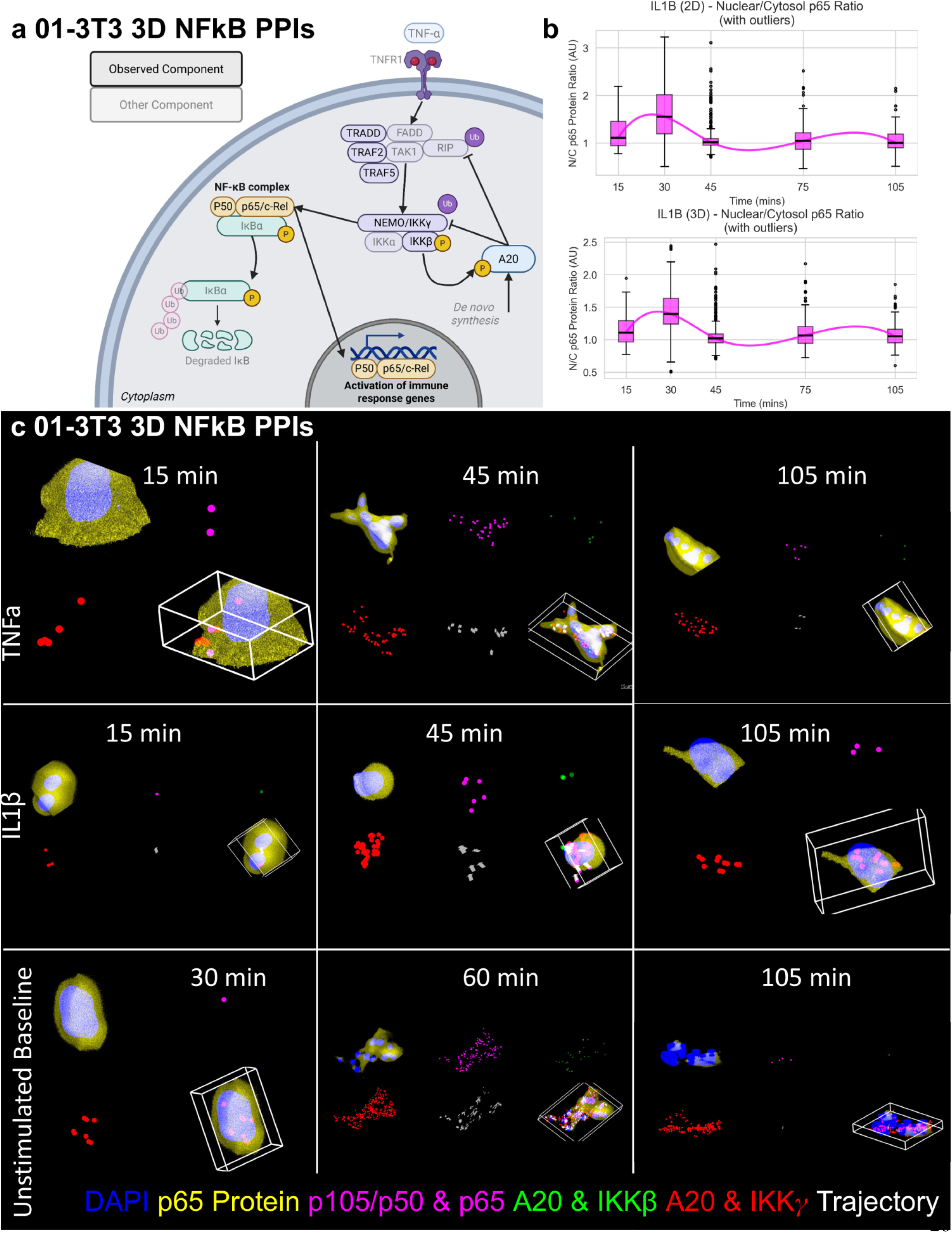
3D NFκB PPI dynamics in TNFα-and IL-1β-stimulated NIH-3T3 mouse fibroblasts. **(a)** Schematic of the observed NFκB signaling components profiled by iseqPLA, depicting canonical pathway members including the IKK complex (IKKα/β/γ), TRADD, TRAF2, TRAF5, RIP1, p105/p50, p65, A20, and downstream transcriptional targets. Created with BioRender.com. **(b)** Nuclear-to-cytoplasmic (N/C) p65 protein ratio over time (15–105 min) after IL-1β stimulation, computed from 2D maximum intensity projections (top) and 3D volumetric reconstructions (bottom). Box plots show the median and interquartile range with individual-cell outliers, and the magenta trend line traces the smoothed mean N/C ratio. The 2D and 3D methods give qualitatively similar translocation kinetics, but 3D lowers cell-to-cell variance and separates nuclear from cytoplasmic signal more cleanly. **(c)** Representative 3D rendered images of single NIH-3T3 fibroblasts at 15, 45, and 105 min post-stimulation for TNFα (top row), IL-1β (middle row), and unstimulated DMSO baseline (bottom row, shown at 30, 60, and 105 min). In total, 2,780 cells were imaged across all timepoints and conditions. Each panel pair shows the cell body rendering alongside a 3D bounding box view. Colors indicate DAPI (blue), p65 protein (yellow), p105/p50 & p65 PLA (magenta), A20 & IKKβ PLA (green), A20 & IKKγ PLA (red), and PPI trajectory (white). Scale bar: indicated per panel.

Fibroblasts (NIH-3T3) were stimulated with TNFα (10 ng/ml), IL-1β (1 ng/ml), or a DMSO vehicle control and fixed at 15-minute intervals across 0–105 minutes, staggered over a single 24-well plate (Supplementary Fig. 1). We chose the IL-1β concentration from values commonly used in fibroblast NFκB activation studies ^[7,41,42]^ and may sit below EC100 for maximal activation. Cell-count heatmaps (Supplementary Fig. 1) confirm adequate sampling in every condition. We did not apply lipopolysaccharide (LPS) to the NIH-3T3 fibroblast experiments, since fibroblasts lack the TLR4/MD-2 co-receptor complex needed to recognize LPS; LPS was instead reserved for the macrophage-and monocyte-containing co-cultures, where the myeloid cells supply the MD-2 cofactor that confers LPS responsiveness.

3D reconstruction of the p65 protein signal showed nuclear enrichment building over time after both TNFα and IL-1β stimulation, peaking at 30–45 minutes and falling off by 105 minutes, the trajectory expected from IκBα resynthesis-driven feedback (Fig. 2c). Over the 45–105 minute window, A20 & IKKβ and A20 & IKKγ PLA signals rose in the stimulated conditions, tracking NFκB-induced A20 expression and its engagement with the IKK complex. Unstimulated cells held low, stable p65 N/C ratios and only sparse PPI puncta throughout. The small baseline fluctuations in those cells likely reflect stochastic basal pathway activity or technical variation inherent to the staggered plate design.

### Cytokines rapidly dissociate upstream NFκB supercomplexes

In NIH-3T3 mouse fibroblasts, both TNFα and IL-1β drove clear dissociation of the upstream TRAF-5_TRADD and TRAF-5_TRAF-2 supercomplexes (representative images in Fig. 3a; population-level quantification in Fig. 3b), a comparison drawn from 1,425 imaged cells. At baseline (0 min) the two PPI signals formed abundant cytoplasmic puncta clusters, as expected for constitutive Complex I. After stimulation, both faded by 45–60 minutes (Fig. 3a), consistent with TRAF2/TRAF5 releasing from TRADD as Complex I is remodeled ^[10]^. Unstimulated cells kept high signal throughout, confirming that dissociation is stimulus-dependent. We define the “p65 nuclear activation score” (Fig. 3b, right) as the mean N/C ratio of p65 fluorescence measured from 3D volumetric segmentation.

**Fig. 3.**
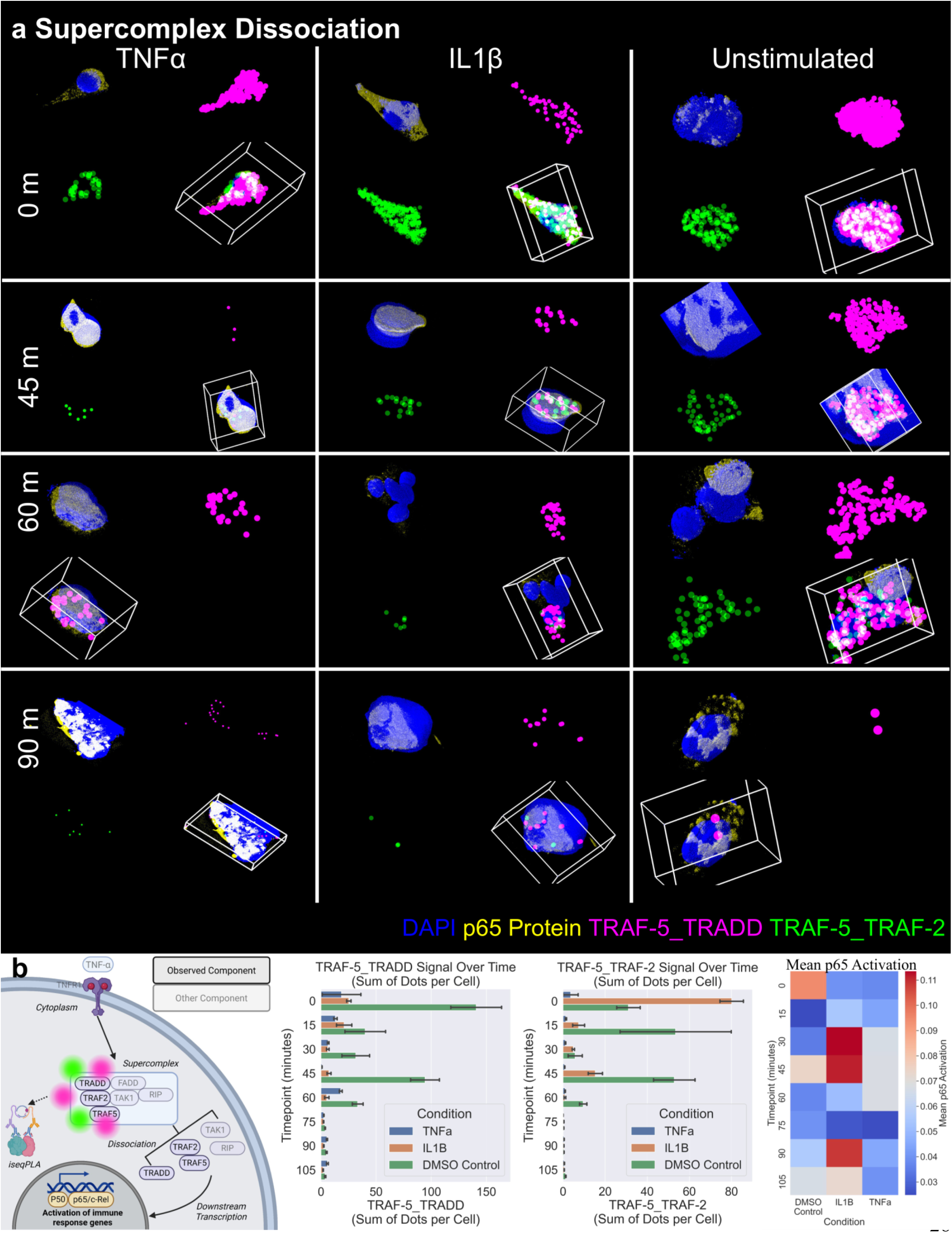
Cytokine-induced dissociation of upstream NFκB supercomplexes in NIH-3T3 mouse fibroblasts. **(a)** Representative 3D rendered images of NIH-3T3 fibroblasts at 0, 45, 60, and 90 min following stimulation with TNFα (10 ng/ml, left column), IL-1β (1 ng/ml, center column), or DMSO unstimulated control (right column). In total, 1,425 cells were imaged across all timepoints and conditions. Each panel shows the cell body and an orthogonal 3D bounding box view. Colors indicate DAPI (blue), p65 protein (yellow), TRAF-5_TRADD PLA (magenta), and TRAF-5_TRAF-2 PLA (green). At early time points (0 min), abundant TRAF-5_TRADD and TRAF-5_TRAF-2 PPI puncta are visible, reflecting intact supercomplex assembly. Following cytokine stimulation, both signals diminish by 45–60 min, consistent with supercomplex dissociation and downstream signal propagation. Unstimulated cells retain elevated PPI puncta throughout the time course. **(b)** Left: schematic of TNFα-induced NFκB supercomplex formation and dissociation, emphasizing how iseqPLA detects the TRAF-5_TRADD and TRAF-5_TRAF-2 interactions at the membrane-proximal Complex I, which then dissociate to drive downstream activation of the p65/p50 transcriptional complex. Center left: TRAF-5_TRADD PPI signal (sum of dots per cell) over time (0–105 min) for TNFα (blue), IL-1β (orange), and DMSO control (green), peaking in unstimulated cells and dropping sharply after cytokine stimulation. Center right: the same quantification for TRAF-5_TRAF-2. Right: heatmap of the mean p65 nuclear activation score (mean nuclear-to-cytoplasmic p65 ratio) across conditions and timepoints, capturing the inverse relationship between upstream supercomplex abundance and downstream p65 nuclear translocation. Bars are means with SEM across cells; per-condition cell counts by timepoint are given in Supplementary Fig. 1a (1,425 segmented cells in total, of which 1,236 carried measurable PLA components and enter the quantification). Connected-component counts for the same plate from the saturation-aware reanalysis are shown in Fig. 4c. Created with BioRender.com.

Counting PPI signal as the sum of RCA puncta per cell reproduced the visual trend (Fig. 3b). Unstimulated cells carried the highest per-cell puncta counts for TRAF-5_TRADD across the early time course, whereas for TRAF-5_TRAF-2 the 0-minute maximum came from the IL-1β well and the unstimulated condition became the highest from 15 minutes onward; the corresponding connected-component counts from the saturation-aware reanalysis are given in Fig. 4c. The spread between conditions at the 0-minute timepoint is well-to-well variability, not biology, since every well was fixed before any cytokine was added. TNFα dissociated the fastest, with TRAF-5_TRADD dropping sharply by 45 minutes, whereas IL-1β gave a blunted response, expected from the distinct receptor-proximal wiring of IL1R, which routes to IKK through IRAK-TRAF6 rather than TRADD-TRAF2/5. The p65 nuclear activation heatmap mirrored this, showing the inverse timing between upstream supercomplex abundance and downstream p65 nuclear enrichment. Error bars represent the standard error of the mean (SEM).

**Fig. 4.**
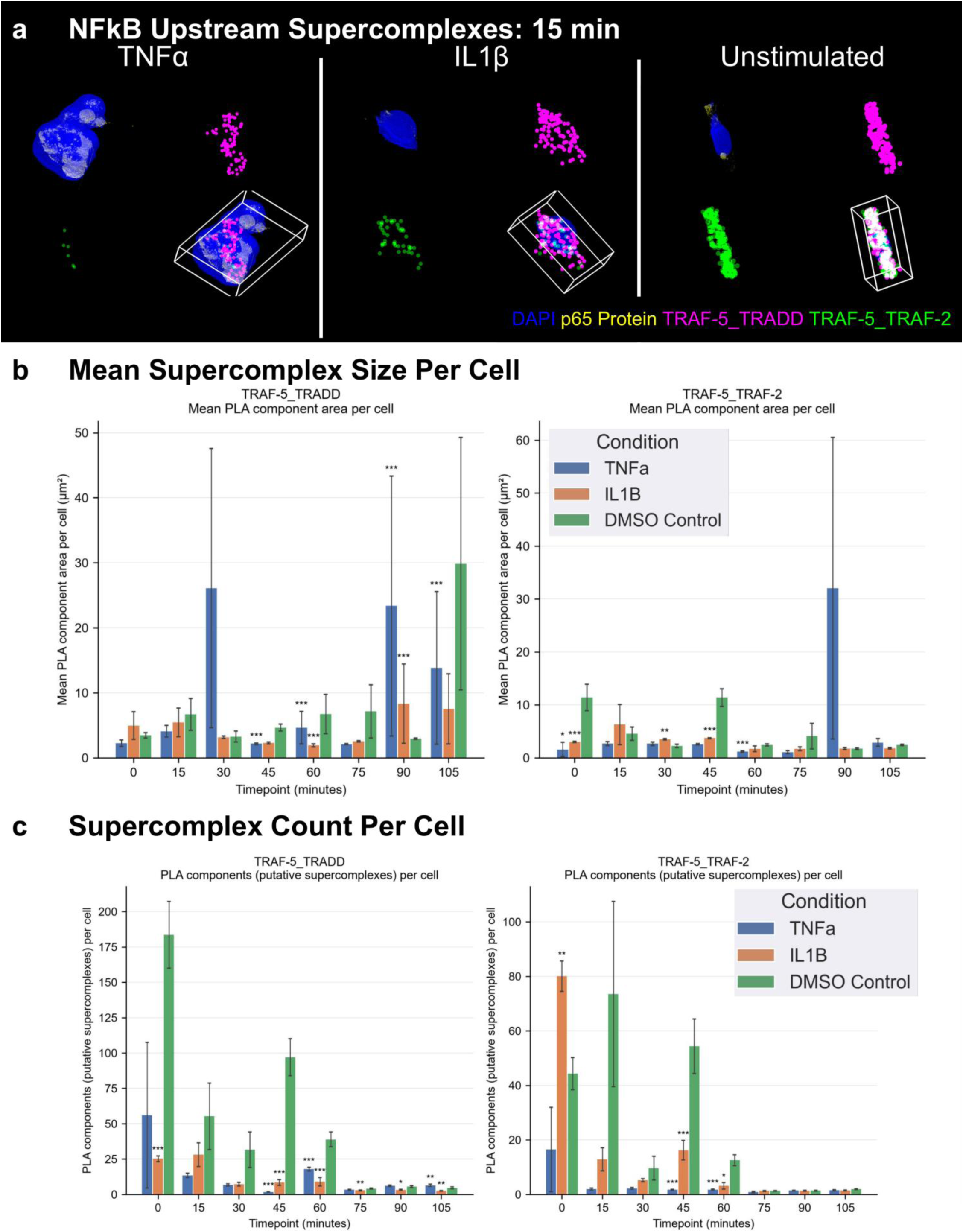
Spatiotemporal quantification of NFκB upstream supercomplex size and count in NIH-3T3 fibroblasts. **(a)** Representative 3D rendered images of NIH-3T3 fibroblasts at 15 min post-stimulation with TNFα (left), IL-1β (center), or DMSO unstimulated control (right), showing DAPI (blue), p65 protein (yellow/magenta), TRAF-5_TRADD PLA (cyan), and TRAF-5_TRAF-2 PLA (magenta/green) with 3D bounding box renderings. At this early time point, supercomplexes remain abundant across all conditions, enabling quantification of their baseline spatial properties prior to dissociation. **(b)** Mean supercomplex area per cell (μm²) over time (0–105 min) for TRAF-5_TRADD (left) and TRAF-5_TRAF-2 (right) PPIs across TNFα (blue), IL-1β (orange), and DMSO control (green) conditions. Mean component area is a morphological descriptor rather than a count: the unstimulated control averages ∼6.7 μm² for TRAF-5_TRADD at 15 min and ∼11.4 μm² for TRAF-5_TRAF-2 at 0 min, while the largest values in the series occur under TNFα at 30, 90 and 105 min (26.1, 23.4 and 13.8 μm² for TRAF-5_TRADD) with standard errors of comparable magnitude (21.5, 20.0 and 11.7 μm²), reflecting a small number of merged, high-density components rather than larger native assemblies (Supplementary Fig. 2). Error bars represent SEM across cells within each condition; per-group n ranges from 2 to 97 cells and is listed by condition and timepoint in Supplementary Fig. 1a. Asterisks denote two-sided Mann–Whitney U tests against the matched DMSO control with Benjamini– Hochberg adjustment within the panel (*p < 0.05, **p < 0.01, ***p < 0.001; ns, not significant). **(c)** Number of supercomplexes per cell over time for TRAF-5_TRADD (left) and TRAF-5_TRAF-2 (right) across the same three conditions. Unstimulated cells exhibit the highest counts at the earliest timepoints (∼184 TRAF-5_TRADD per cell at 0 min and ∼74 TRAF-5_TRAF-2 per cell at 15 min; the 0-minute TRAF-5_TRAF-2 maximum of ∼80 per cell comes from the IL-1β well), with a rapid decline following cytokine stimulation. Error bars represent SEM across cells within each condition, with per-group n and statistics as in (b). Figs. 4b and 4c present complementary quantifications (mean cluster area and cluster count, respectively) derived from the same imaging datasets.

To capture supercomplex morphology, we tracked it across the full time course (Fig. 4a–c), with representative images at 15 minutes (Fig. 4a). Here “supercomplex area” is the projected area of contiguous RCA signal clusters within a 2D MIP, and “supercomplex count” is the number of discrete clusters per cell from intensity thresholding and connected-component analysis. Because individual RCA products can merge into confluent signal at high interaction density, connected-component counts undercount proximity events in the saturated regime; we therefore treat these clusters as putative supercomplexes and treat supercomplex count and area as morphological descriptors of proximity signal, not absolute molecular counts. Single-punctum reference areas estimated per probe pair from the area distributions ranged from ≈45 to 207 px² depending on the probe pair, and the fraction of PLA signal residing in merged components larger than 3× this reference rises from ∼0% in low-signal cells to 81–98% in the densest cells, while stimulation increases primarily the number of components per cell, not their size (analysis of plates 004A–C; Supplementary Fig. 2). Component counts were highest in unstimulated cells at the earliest timepoints (∼184 TRAF-5_TRADD per cell at 0 min, falling to ∼55 by 15 min; ∼74 TRAF-5_TRAF-2 per cell at 15 min, with the 0-minute TRAF-5_TRAF-2 maximum of ∼80 per cell coming from the IL-1β well; Fig. 4c). TNFα produced the steepest reductions in count, collapsing TRAF-5_TRAF-2 to ∼2 components per cell by 45–60 minutes. Mean component area did not track count (Fig. 4b): the unstimulated control averaged ∼6.7 μm² for TRAF-5_TRADD at 15 min and ∼11.4 μm² for TRAF-5_TRAF-2 at 0 min, while the largest values in the series belong to TNFα at 30, 90 and 105 min (26.1, 23.4 and 13.8 μm² for TRAF-5_TRADD) and carry standard errors of comparable magnitude (21.5, 20.0 and 11.7 μm²). Those excursions come from a small number of merged, high-density components rather than from larger native assemblies, the saturation behaviour characterized in Supplementary Fig. 2, so we read component count rather than mean component area as the more reliable descriptor of proximity-signal abundance. Error bars represent SEM.

### ECM coating dictates the NFκB signaling response

Across three substrate coatings, NIH-3T3 fibroblast p65 nuclear translocation at the 45-minute activation peak diverged sharply (Supplementary Fig. 3). Under TNFα, collagen I gave the largest activated fraction (86.8%) and poly-L-lysine an intermediate one (67.2%), while Matrigel held activation down to 18.2% (all pairwise comparisons p < 0.001). Under IL-1β, collagen I (87.5%) and poly-L-lysine (87.6%) were essentially matched (ns), and Matrigel again lagged (29.8%). DMSO baselines stayed low on every coating (1.6–15.4%).

We applied the three substrates at their standard working concentrations (0.01% poly-L-lysine, 50 μg/ml collagen I, 500 μg/ml Matrigel) instead of matching them by molarity, so teasing substrate identity apart from concentration would require a dedicated dose-response series. The suppression seen with Matrigel could stem from cytokine sequestration in the dense matrix, altered integrin signaling, or weaker receptor engagement, while the boost from collagen I fits integrin-FAK-IKK crosstalk that amplifies TNFα-induced p65 translocation ^[33]^. ECM coating is therefore a decisive experimental variable in NFκB studies, and we used poly-L-lysine and collagen I for all subsequent work. A small Matrigel titration (Supplementary Fig. 4) reinforced the point: pushing TNFα or IL-1β higher did not relieve the Matrigel-imposed block on p65 nuclear translocation.

### Macrophage-fibroblast co-cultures reveal paracrine NFκB activation dynamics

We compared NFκB p65 activation across three parallel settings: CCL2-exposed macrophage co-cultures (control), CFASN (CF airway supernatant)-exposed macrophage co-cultures (CF disease), and IMR-90 fibroblast mono-cultures (Supplementary Fig. 1; Figs. 5–6; Supplementary Fig. 5). The CCL2 and CFASN macrophages came from the Tirouvanziam laboratory’s organotypic airway transmigration model and were plated with IMR-90 fibroblasts at a 1:10 ratio. Each co-culture was stimulated with LPS, TNFα, IL-1β, or DMSO for 0–480 minutes and imaged for Phospho-S536 p65, total p65, and Phalloidin.

**Fig. 5.**
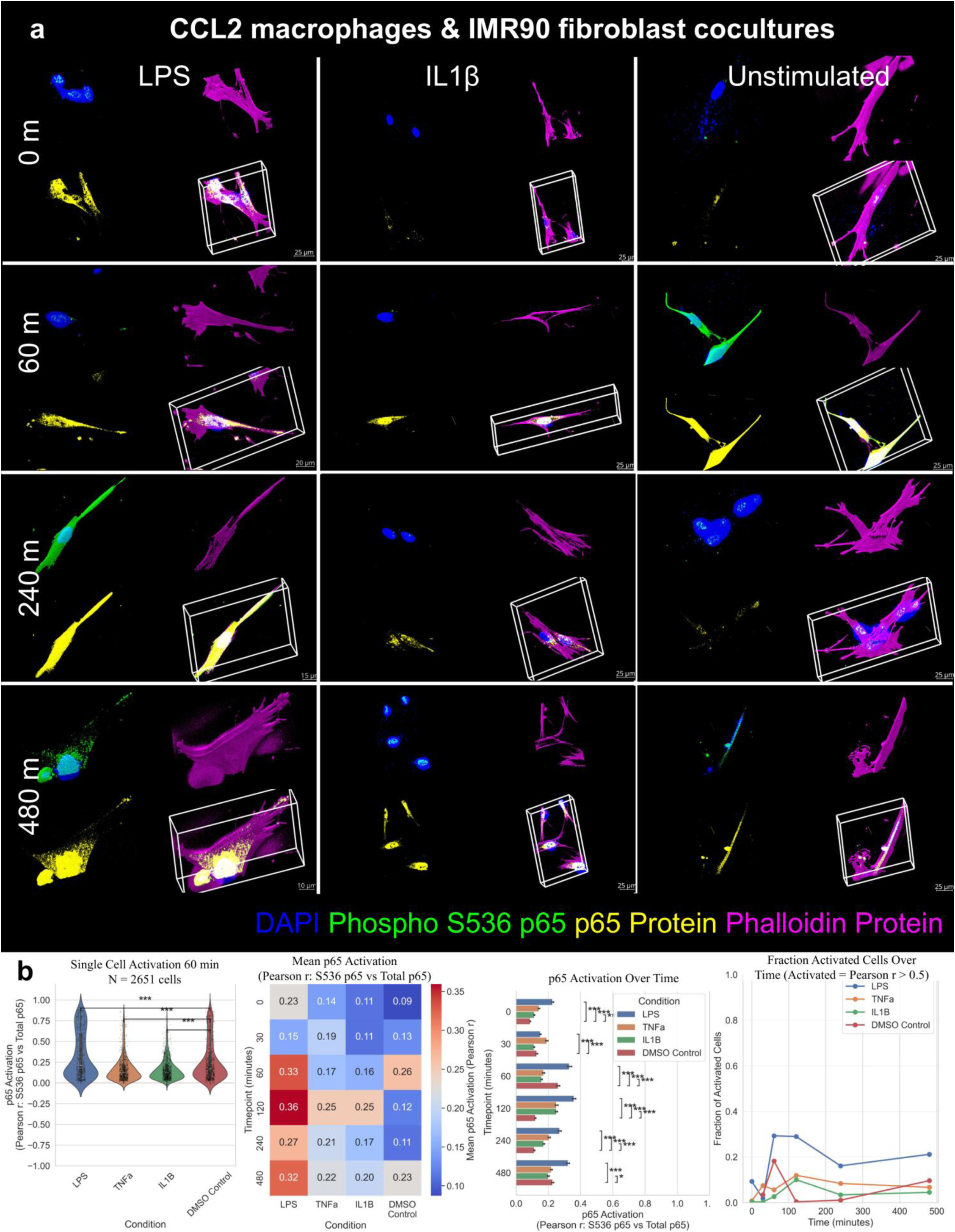
NFκB p65 activation dynamics in CCL2 control macrophage and IMR-90 fibroblast co-cultures. **(a)** Representative 3D rendered images of CCL2-exposed macrophage (control, chemokine attractant only) and IMR-90 fibroblast co-cultures at 0, 60, 240, and 480 min post-stimulation with LPS (10 ng/ml, left column), IL-1β (1 ng/ml, center column), or DMSO unstimulated control (right column). In total, 16,961 cells were imaged across all timepoints and conditions. Colors indicate DAPI (blue), Phospho-S536 p65 (green), total p65 protein (yellow), and Phalloidin (magenta, actin cytoskeleton). The elongated fibroblast morphology and rounder macrophage morphology are distinguishable in 3D reconstructions. Scale bars as indicated per panel. **(b)** Quantitative analyses of single-cell p65 activation (Pearson r of Phospho-S536 p65 vs. total p65) in CCL2 co-cultures. Far left: Violin plots of p65 activation scores at 60 min across LPS, TNFα, IL-1β, and DMSO control conditions with statistical significance annotations. Center left: Heatmap of mean p65 activation scores across all four stimulation conditions (rows) and timepoints 0–480 min (columns). Center right: Mean p65 activation over time for all conditions with error bars and pairwise statistical comparisons. Far right: Fraction of p65-activated cells (Pearson r > 0.5 threshold) over time, demonstrating LPS-and TNFα-stimulated CCL2 co-cultures achieving the highest activation fractions at early timepoints, followed by gradual resolution consistent with canonical NFκB feedback termination. The 60-minute panel comprises n = 2,651 cells (LPS 481, TNFα 692, IL-1β 798, DMSO 680) and error bars are SEM across cells; per-condition counts for every timepoint are given in Supplementary Fig. 1b. Asterisks denote two-sided Mann–Whitney U tests against the matched DMSO control with Benjamini– Hochberg adjustment within the panel (*p < 0.05, **p < 0.01, ***p < 0.001; ns, not significant).

**Fig. 6.**
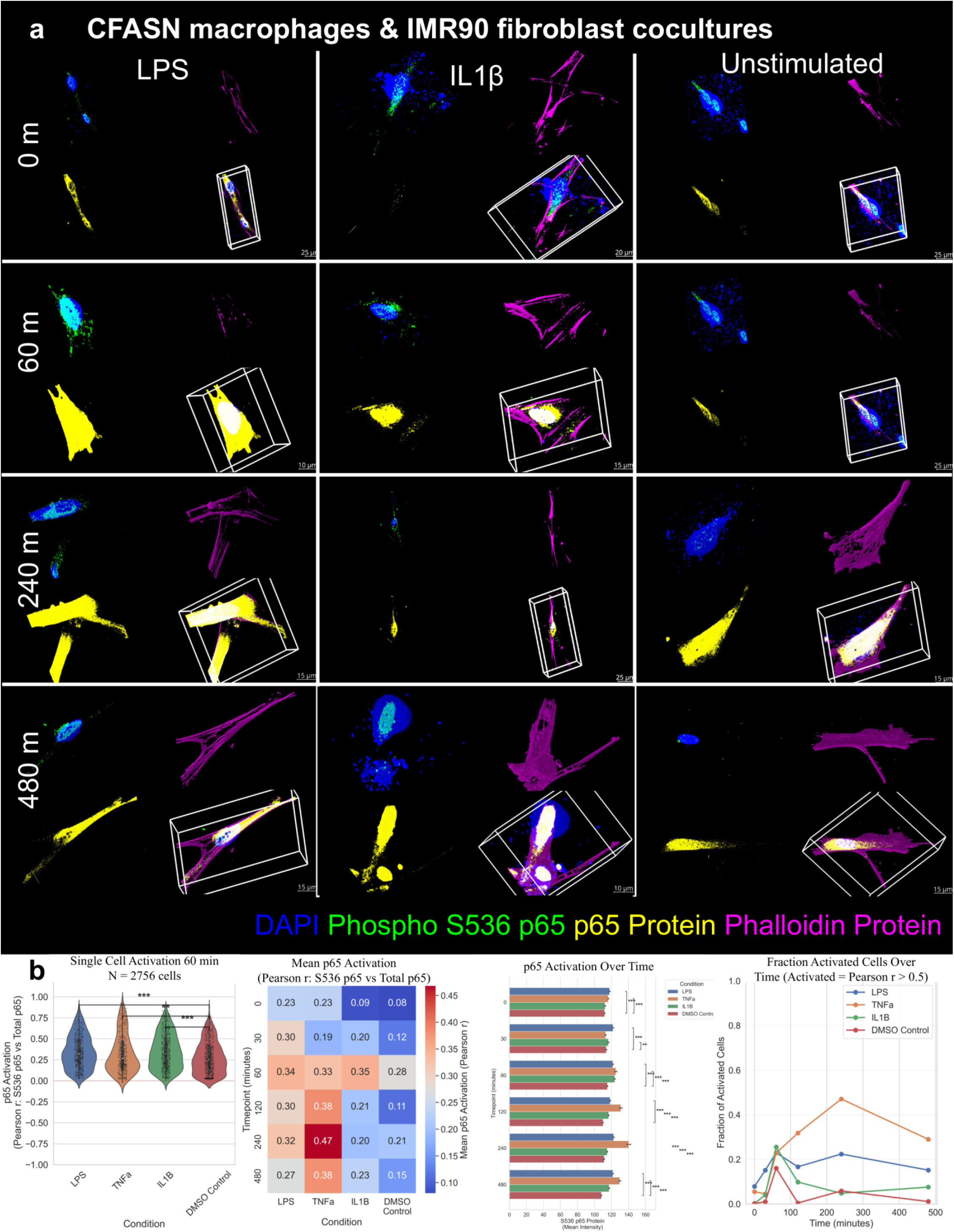
NFκB p65 activation dynamics in CF airway supernatant-exposed (CFASN) macrophage and IMR-90 fibroblast co-cultures. **(a)** Representative 3D rendered images of CFASN macrophage (cystic fibrosis patient-derived, transmigrated macrophages exposed to CF airway supernatant) and IMR-90 fibroblast co-cultures at 0, 60, 240, and 480 min following stimulation with LPS (10 ng/ml, left column), IL-1β (1 ng/ml, center column), or DMSO unstimulated control (right column). In total, 15,617 cells were imaged across all timepoints and conditions. Colors indicate DAPI (blue), Phospho-S536 p65 (green), total p65 protein (yellow), and Phalloidin (magenta). Compared to CCL2 control macrophage co-cultures (Fig. 5), CFASN co-cultures display altered p65 activation patterns across stimulation conditions, reflecting the hyperinflammatory phenotype of CF-derived transmigrated macrophages. Scale bars as indicated per panel. **(b)** Quantitative analyses of single-cell p65 activation. Far left: Violin plots of p65 activation (Pearson r Phospho-S536 p65 vs. total p65) at 60 min across conditions, with statistical significance markers; CFASN co-cultures exhibit elevated baseline and stimulated activation compared to CCL2 controls. Center left: Heatmap of mean p65 activation scores across conditions and timepoints (0–480 min); TNFα-stimulated CFASN co-cultures showed the highest mean activation score (0.47 at 240 min), while IL-1β-stimulated co-cultures showed elevated activation relative to CCL2 counterparts (0.35 versus 0.16 for CCL2 IL-1β at 60 min), consistent with dysregulated NFκB signaling in the CF inflammatory context. Center right: Mean phospho-S536 p65 intensity per cell over time for all four stimulation conditions, with SEM error bars and pairwise significance annotations; the corresponding Pearson-r activation values are those shown in the heatmap at center left. Far right: Fraction of activated cells over time; TNFα reaches the largest activated fraction of any condition (∼0.47 at 240 min) and LPS shows a secondary peak of ∼0.23 at 240 min before falling to ∼0.15 by 480 min. The 60-minute panel comprises n = 2,756 cells (LPS 622, TNFα 577, IL-1β 791, DMSO 766); per-condition counts for every timepoint are given in Supplementary Fig. 1b. Asterisks denote two-sided Mann–Whitney U tests against the matched DMSO control with Benjamini–Hochberg adjustment within the panel (*p < 0.05, **p < 0.01, ***p < 0.001; ns, not significant).

Here we relied on Phospho-S536 p65 rather than iseqPLA, simply because a ∼24-hour-per-cycle iseqPLA workflow was impractical across 72 conditions. Activation was scored per cell as the Pearson correlation (r) between Phospho-S536 p65 and total p65 intensity, and cells with r > 0.5 were called “activated”. This whole-cell Phospho-S536/total-p65 correlation is a co-localization proxy for pathway engagement rather than a direct measure of nuclear translocation; a direct per-cell nuclear-to-cytoplasmic (N/C) quantification of the phospho-p65 pool is reported alongside it in Supplementary Fig. 6, where the N/C ratio does not separate the stimulation conditions that the correlation metric resolves. No acquisition saturation was detected in the p65 channels (recorded maxima of 0.9×10⁴ to 4.0×10⁴ out of 65,535 16-bit levels across all 5,407 cells, and no cell exceeded 1% saturated voxels), the near-saturated appearance of p65 signal in figure images reflects display-contrast scaling rather than detector saturation, so we interpret the activation scores as relative, within-experiment comparisons.

In the CCL2 co-cultures (representative images in Fig. 5a; 16,961 cells imaged), LPS drove the strongest and fastest response, with the activated fraction climbing to ∼0.3 by 60 minutes before easing off (population-level quantification in Fig. 5b). At 60 minutes TNFα and IL-1β sat at or below the DMSO baseline in the CCL2 co-cultures (mean activation 0.17 and 0.16 versus 0.26), although both rose above it at 120 and 240 minutes (0.25 and 0.25 versus 0.12 at 120 min), and DMSO controls stayed low throughout.

In the CFASN co-cultures (representative images in Fig. 6a; 15,617 cells imaged), the dynamics shifted in a way that fits CF macrophage hyperinflammation. TNFα-stimulated CFASN co-cultures reached the highest mean activation score (0.47 at 240 min), and IL-1β stimulation raised activation above the CCL2 comparators (0.35 vs. 0.16 at 60 min; population-level quantification in Fig. 6b). At 60 minutes, violin-plot comparisons confirmed the elevations were statistically significant, pointing to CF airway conditioning as an amplifier of paracrine NFκB signaling in this preliminary, single-donor model. The LPS-stimulated activated fraction rose again to a secondary peak of ∼0.23 at 240 minutes before falling to ∼0.15 by 480 minutes, while TNFα produced the largest activated fraction of any condition (∼0.47 at 240 minutes; Fig. 6b). The imaged totals (16,961 cells for Fig. 5 and 15,617 cells for Fig. 6) span all four stimulations and six timepoints, whereas the Fig. 5b and 6b quantification panels display the 60-minute subset (n = 2,651 and n = 2,756 cells, respectively), matching the per-condition cell-count heatmaps in Supplementary Fig. 1b; essentially no cells are discarded by quality-control filtering (<1%, only degenerate segmentations).

By contrast, IMR-90 mono-cultures (Supplementary Fig. 5; 13,408 cells) responded only weakly to LPS, as expected when fibroblasts lack the macrophage-provided MD-2 cofactor. IL-1β pushed activation steadily upward through 480 minutes, yet the fraction crossing r > 0.5 stayed low, reaching only ∼0.12 by 480 minutes against the ∼0.3 the macrophage co-cultures reached by 60 minutes.

### Monocyte-fibroblast co-cultures extend supercomplex profiling to disease-relevant immune microenvironments

To add spatial supercomplex information on top of the phospho-p65 readouts, we ran 3D iseqPLA on the A20 & IKKβ and TRAF-5 & TRAF-2 PPIs in monocyte-fibroblast co-cultures from a separate donor cohort (courtesy of Dr. Kopp’s laboratory). Healthy and CF patient-derived monocytes were each combined with IMR-90 fibroblasts at a 1:10 ratio, alongside fibroblast mono-culture controls, and all conditions were stimulated with LPS, TNFα, IL-1β, or DMSO for 0–480 minutes.

In the healthy-donor monocyte co-cultures (Fig. 7a; 10,570 cells; Supplementary Fig. 7a), both PPI signals were already present as discrete cytoplasmic clusters at baseline and then remodeled over time once stimulated. Both supercomplex count (Fig. 7b,c) and area (Fig. 7d,e) moved in a condition-dependent way, with cytokine-stimulated wells shifting in abundance and morphology relative to controls.

**Fig. 7.**
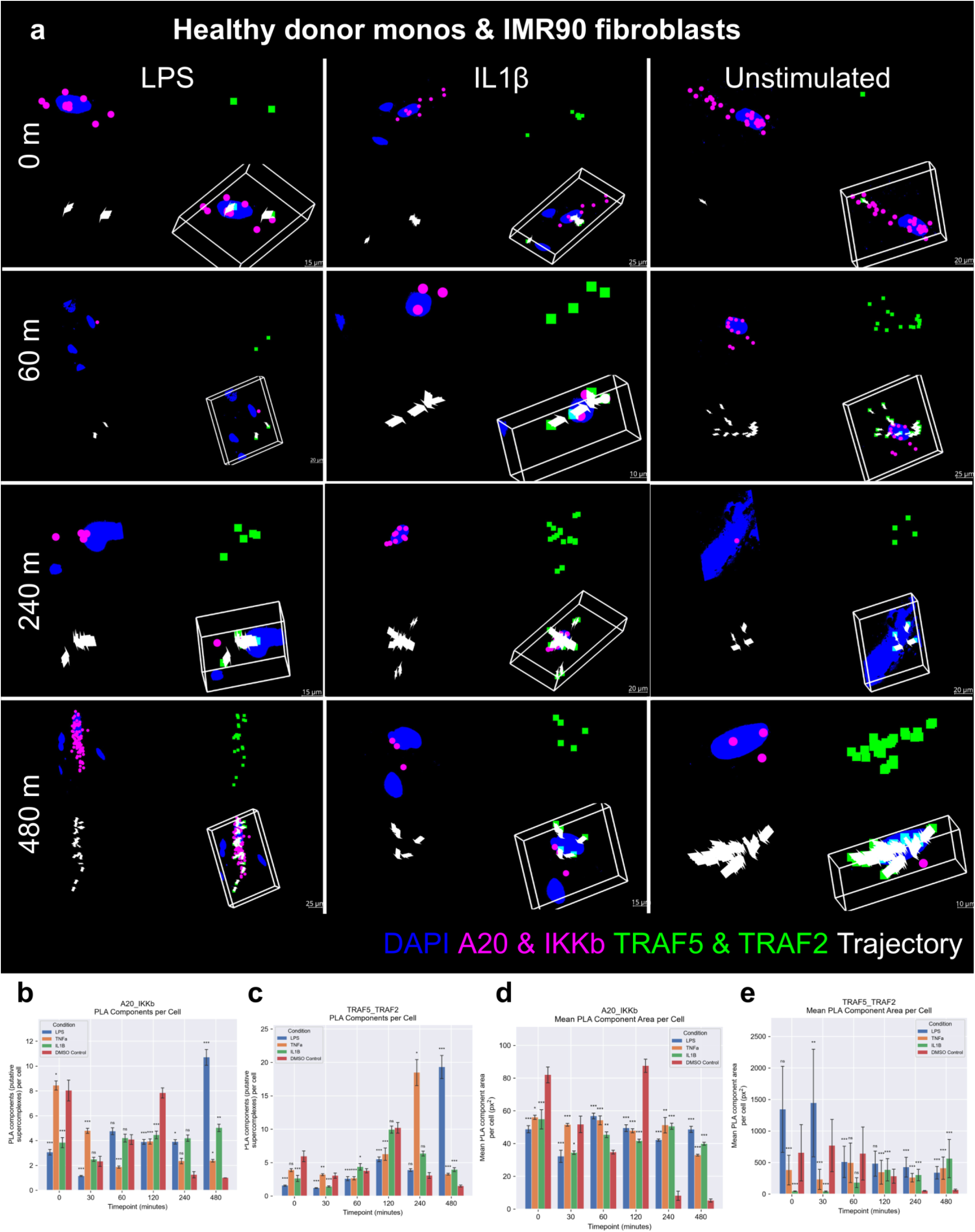
NFκB supercomplex dynamics in healthy donor monocytes and IMR-90 fibroblast co-cultures. **(a)** Representative 3D rendered images of healthy donor-derived monocyte and IMR-90 fibroblast co-cultures at 0, 60, 240, and 480 min post-stimulation with LPS (10 ng/ml, left column), IL-1β (1 ng/ml, center column), or DMSO unstimulated control (right column). In total, 10,570 cells were imaged across all timepoints and conditions; cell count heatmaps are presented in Supplementary Fig. 7a. Colors indicate DAPI (blue), A20 & IKKβ PLA (magenta), and TRAF-5 & TRAF-2 PLA (green). Each panel shows the cell body rendering alongside a 3D bounding box view. Scale bars as indicated per panel. **(b)** Number of A20_IKKβ supercomplexes per cell over time (0–480 min) for LPS (blue), TNFα (orange), IL-1β (green), and DMSO control (red) conditions. Error bars represent SEM. **(c)** Number of TRAF-5_TRAF-2 supercomplexes per cell over time across the same four conditions. Error bars represent SEM. **(d)** Mean A20_IKKβ component area per cell (px²; 1 px² ≈ 0.0125 μm² at the acquisition pixel size) over time across all four stimulation conditions. Error bars represent SEM. **(e) Mean TRAF-5_TRAF-2 component area per cell (px²; 1 px² ≈ 0.0125 μm² at the acquisition pixel size) over time across all four stimulation conditions. Error bars represent SEM. Panels (b–e) draw on 6,497 cell-marker measurements for A20_IKKβ and 6,021 for TRAF-5_TRAF-2 (per-group n = 4–526 and 15– 463 cells, respectively). Asterisks denote two-sided Mann–Whitney U tests against the matched DMSO control with Benjamini–Hochberg adjustment within the panel (*p < 0.05, **p < 0.01, ***p < 0.001; ns, not significant).**

The CF patient-derived monocyte co-cultures (Fig. 8a; 10,984 cells; Supplementary Fig. 7b) followed a broadly similar pattern, but the quantification exposed differences that fit dysregulated NFκB signaling. TRAF-5 & TRAF-2 supercomplexes were more abundant at later timepoints under certain conditions than in the healthy co-cultures (Fig. 8c), hinting that CF monocyte-derived paracrine signals reshape upstream supercomplex remodeling, and the mean supercomplex areas (Fig. 8d,e) differed in a condition-specific manner.

**Fig. 8.**
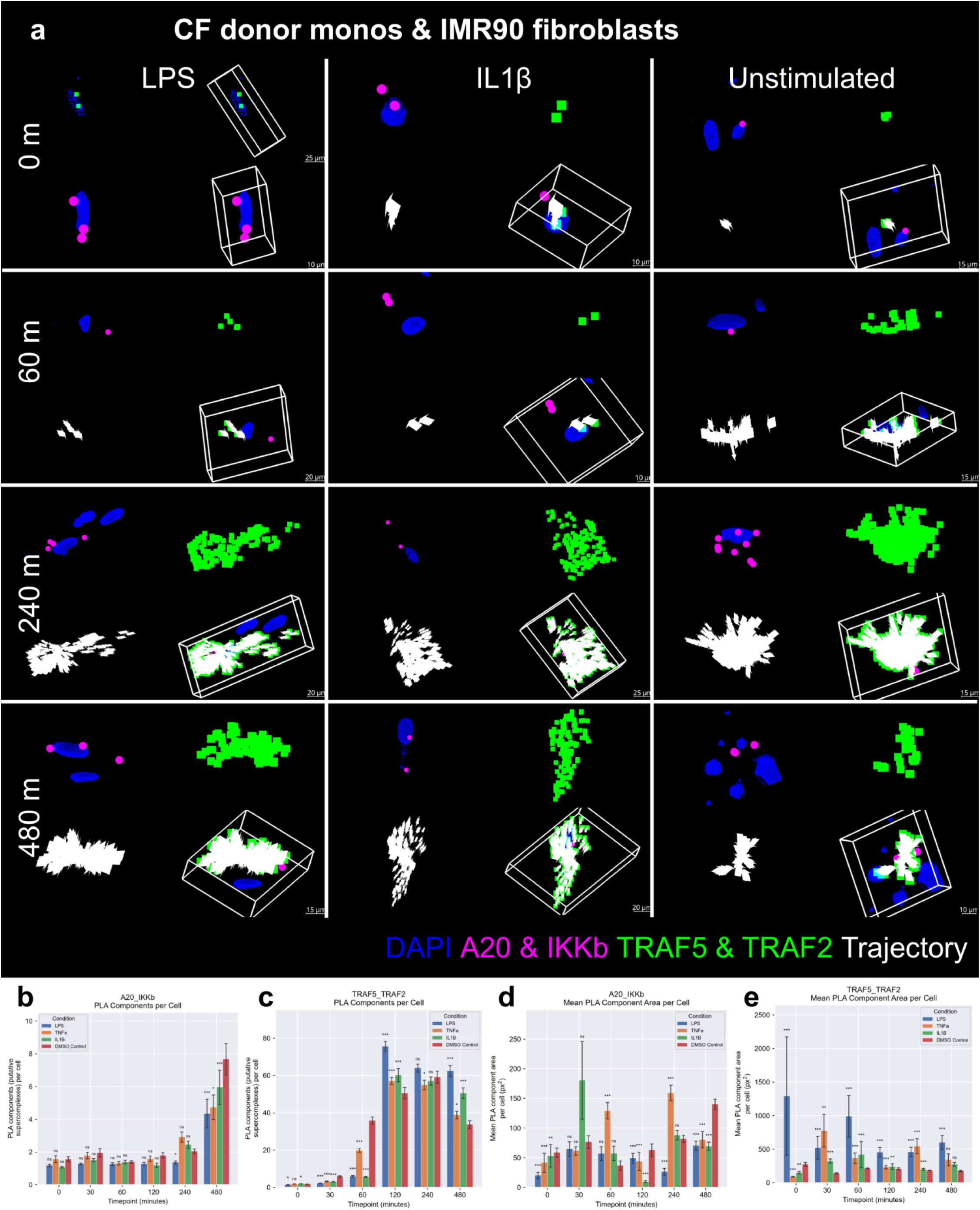
NFκB supercomplex dynamics in CF patient-derived monocyte and IMR-90 fibroblast co-cultures. **(a)** Representative 3D rendered images of CF patient-derived monocyte and IMR-90 fibroblast co-cultures at 0, 60, 240, and 480 min post-stimulation with LPS (10 ng/ml, left column), IL-1β (1 ng/ml, center column), or DMSO unstimulated control (right column). In total, 10,984 cells were imaged across all timepoints and conditions; cell count heatmaps are presented in Supplementary Fig. 7b. Colors indicate DAPI (blue), A20 & IKKβ PLA (magenta), and TRAF-5 & TRAF-2 PLA (green). Each panel shows the cell body rendering alongside a 3D bounding box view. Compared to healthy monocyte co-cultures (Fig. 7), CF monocyte co-cultures display altered supercomplex dynamics consistent with dysregulated NFκB signaling in the CF inflammatory context. Scale bars as indicated per panel. **(b)** Number of A20_IKKβ supercomplexes per cell over time (0–480 min) for LPS (blue), TNFα (orange), IL-1β (green), and DMSO control (red) conditions. Error bars represent SEM. **(c)** Number of TRAF-5_TRAF-2 supercomplexes per cell over time across the same four conditions. Error bars represent SEM. **(d)** Mean A20_IKKβ component area per cell (px²; 1 px² ≈ 0.0125 μm² at the acquisition pixel size) over time across all four stimulation conditions. Error bars represent SEM. **(e) Mean TRAF-5_TRAF-2 component area per cell (px²; 1 px² ≈ 0.0125 μm² at the acquisition pixel size) over time across all four stimulation conditions. Error bars represent SEM. Panels (b–e) draw on 1,860 cell-marker measurements for A20_IKKβ and 8,423 for TRAF-5_TRAF-2 (per-group n = 16–175 and 100–623 cells, respectively). Asterisks denote two-sided Mann–Whitney U tests against the matched DMSO control with Benjamini–Hochberg adjustment within the panel (*p < 0.05, **p < 0.01, ***p < 0.001; ns, not significant).**

IMR-90 mono-cultures (Fig. 9a; 7,719 cells; Supplementary Fig. 7c) remodeled little under LPS relative to either co-culture, again consistent with fibroblasts being LPS-insensitive without monocyte-derived signals. IL-1β elicited the sharpest changes, as expected for direct IL1R engagement. Read together, the healthy, CF, and mono-culture arms indicate that disease-associated immune conditioning tunes both the abundance and the morphology of NFκB PPI assemblies.

**Fig. 9.**
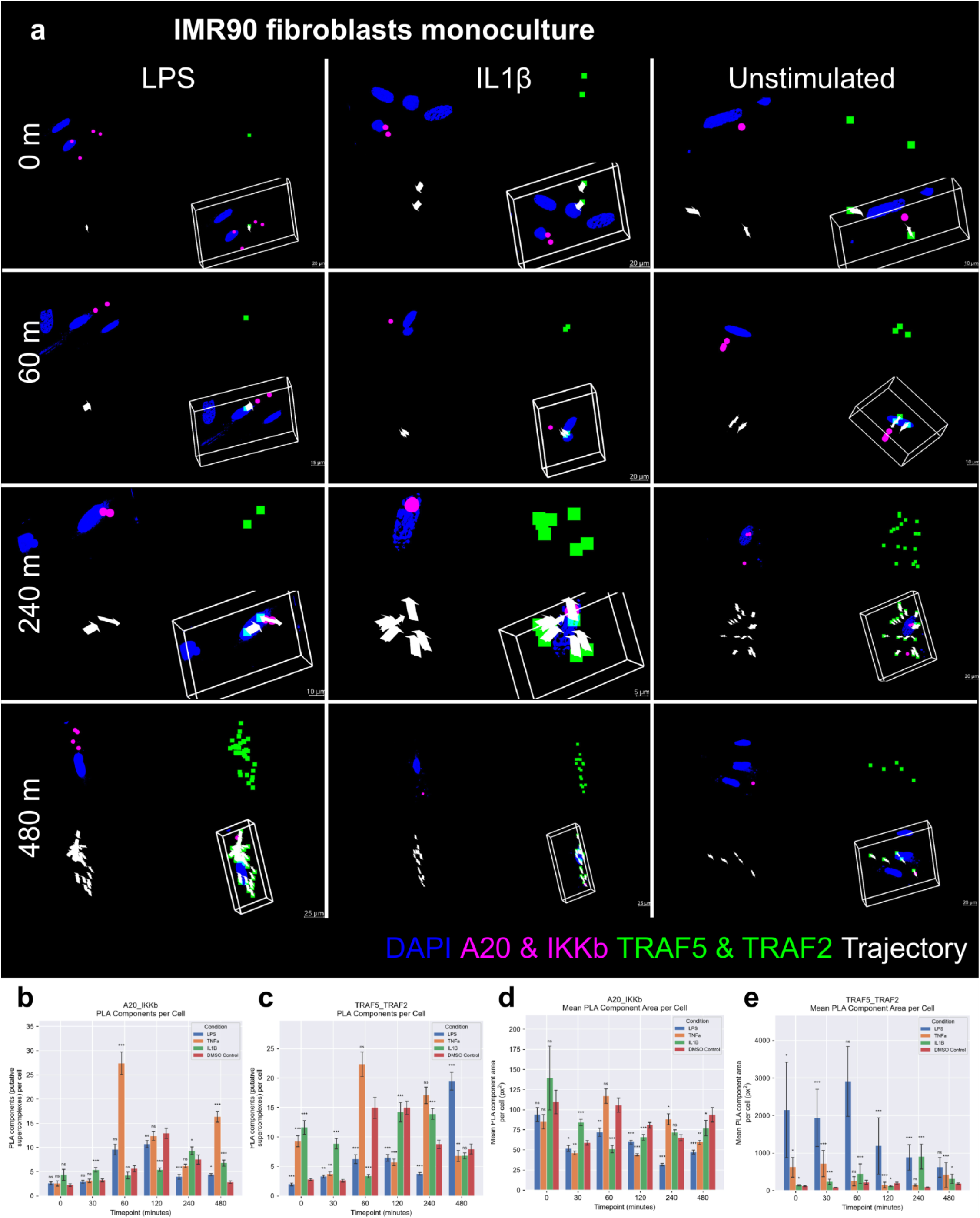
NFκB supercomplex dynamics in corresponding IMR-90 fibroblast mono-cultures. **(a)** Representative 3D rendered images of IMR-90 fibroblast mono-cultures (no monocytes) at 0, 60, 240, and 480 min post-stimulation with LPS (10 ng/ml, left column), IL-1β (1 ng/ml, center column), or DMSO unstimulated control (right column). In total, 7,719 cells were imaged across all timepoints and conditions; cell count heatmaps are presented in Supplementary Fig. 7c. Colors indicate DAPI (blue), A20 & IKKβ PLA (magenta), and TRAF-5 & TRAF-2 PLA (green). Compared to healthy (Fig. 7) and CF (Fig. 8) monocyte co-culture conditions, fibroblast mono-cultures show attenuated supercomplex remodeling, particularly under LPS stimulation, consistent with the known LPS insensitivity of fibroblasts in the absence of paracrine monocyte-derived signaling. Scale bars as indicated per panel. **(b)** Number of A20_IKKβ supercomplexes per cell over time (0–480 min) for LPS (blue), TNFα (orange), IL-1β (green), and DMSO control (red) conditions. Error bars represent SEM. **(c)** Number of TRAF-5_TRAF-2 supercomplexes per cell over time across the same four conditions. Error bars represent SEM. **(d)** Mean A20_IKKβ component area per cell (px²; 1 px² ≈ 0.0125 μm² at the acquisition pixel size) over time across all four stimulation conditions. Error bars represent SEM. **(e) Mean TRAF-5_TRAF-2 component area per cell (px²; 1 px² ≈ 0.0125 μm² at the acquisition pixel size) over time across all four stimulation conditions. Error bars represent SEM.** Figs. 9b–e present complementary quantifications (supercomplex count and mean cluster area for A20_IKKβ and TRAF-5_TRAF-2, respectively) derived from the same imaging datasets, paralleling the analyses in Figs. 7b–e and 8b–e. Error bars represent SEM. Panels (b–e) draw on 4,010 cell-marker measurements for A20_IKKβ and 5,680 for TRAF-5_TRAF-2 (per-group n = 47–310 and 143–345 cells, respectively). Asterisks denote two-sided Mann–Whitney U tests against the matched DMSO control with Benjamini–Hochberg adjustment within the panel (*p < 0.05, **p < 0.01, ***p < 0.001; ns, not significant).

### Exploratory scGPT embedding analysis of the NFκB gene panel

To ask, computationally, whether the iseqPLA panel targets are biologically well chosen, we turned to scGPT ^[40]^, a transformer pretrained on large-scale single-cell transcriptomic atlases (Supplementary Fig. 8). Our panel covers only a small set of proteins (TRAF-5, TRAF-2, TRADD, A20, p50/p105, IKKβ, IKKγ, and p65), so we asked whether the genes encoding them fall within inflammation-relevant regions of transcriptional feature space. The intent here is to support the panel design with independent, exploratory evidence, not to uncover new biology.

We fine-tuned scGPT on a curated multi-species in vitro corpus of five publicly available datasets (31,915 cells pre-QC, 27,529 cells post-QC; Supplementary Table 1) to classify control versus inflamed/treated states. The corpus paired four curated mouse in vitro NFκB-relevant datasets (RAW264.7 macrophages, GSE94383; HSPCs/GMPs, GSE199404; 3T3 fibroblast–macrophage co-cultures, GSE189062; MEFs with RelA knockout and complementation, GSE132791; total n = 1,126 pre-QC) with human PBMCs (GSE226488; n = 30,789 pre-QC). For mixed-species training we built a shared gene vocabulary by case-insensitive intersection of mouse and human gene symbols; an alternative MGI (Mouse Genome Informatics)-ortholog-based intersection gave equivalent results. Before training we also removed three technical-artifact gene families using HGNC (HUGO Gene Nomenclature Committee)-curated allowlists (mitochondrial transcripts (MT-prefix), cytoplasmic ribosomal proteins (HGNC group 1054, 90 genes), and canonical hemoglobin subunits (HGNC group 940, 10 genes)) because such genes otherwise dominate feature-importance rankings in fine-tuned single-cell transformers, reflecting dissociation and metabolic stress, not pathway-specific biology. That left a final canonical vocabulary of roughly 13,109 shared gene symbols. Quality control followed the single-cell best-practices recipe: 5-MAD (median absolute deviation) outlier filtering on library size, gene count, and percent-top-20 metrics; a 3-MAD plus 20% absolute ceiling on percent mitochondrial reads; and a 50-gene-per-cell floor, each computed per dataset so the majority human dataset could not set the thresholds for everyone.

A UMAP of the fine-tuned embeddings cleanly separated cells by dataset of origin, perturbation condition, and phenotype (Fig. 10a–c): the human PBMC dataset formed the dominant manifold, while the smaller mouse cell-line datasets (3T3 fibroblasts, HSPCs, MEFs, RAW264.7 macrophages) resolved into distinct sub-populations. Consensus ranking across gradient, SHAP, and perturbation attribution put PLCG2, SDC2, LMNA, CCL5, SRSF5, S100A11, ANXA1, PASK, ZFP36L1, and CDC42 at the head of the top 30 discriminative genes (Fig. 10d). That list leaned heavily toward canonical NFκB-responsive and inflammatory modulators (CCL5, CXCL2, ANXA1, ZFP36L1, ZFP36, CEBPA, IL1RN, AIF1, IFI30) alongside extracellular matrix and fibroblast-identity genes (COL1A1, COL1A2, FBLN2, TAGLN), unsurprising given the corpus, and, as intended, no mitochondrial, ribosomal, or hemoglobin genes surfaced in the top 30, a sign the upstream artifact filtering worked.

**Fig. 10.**
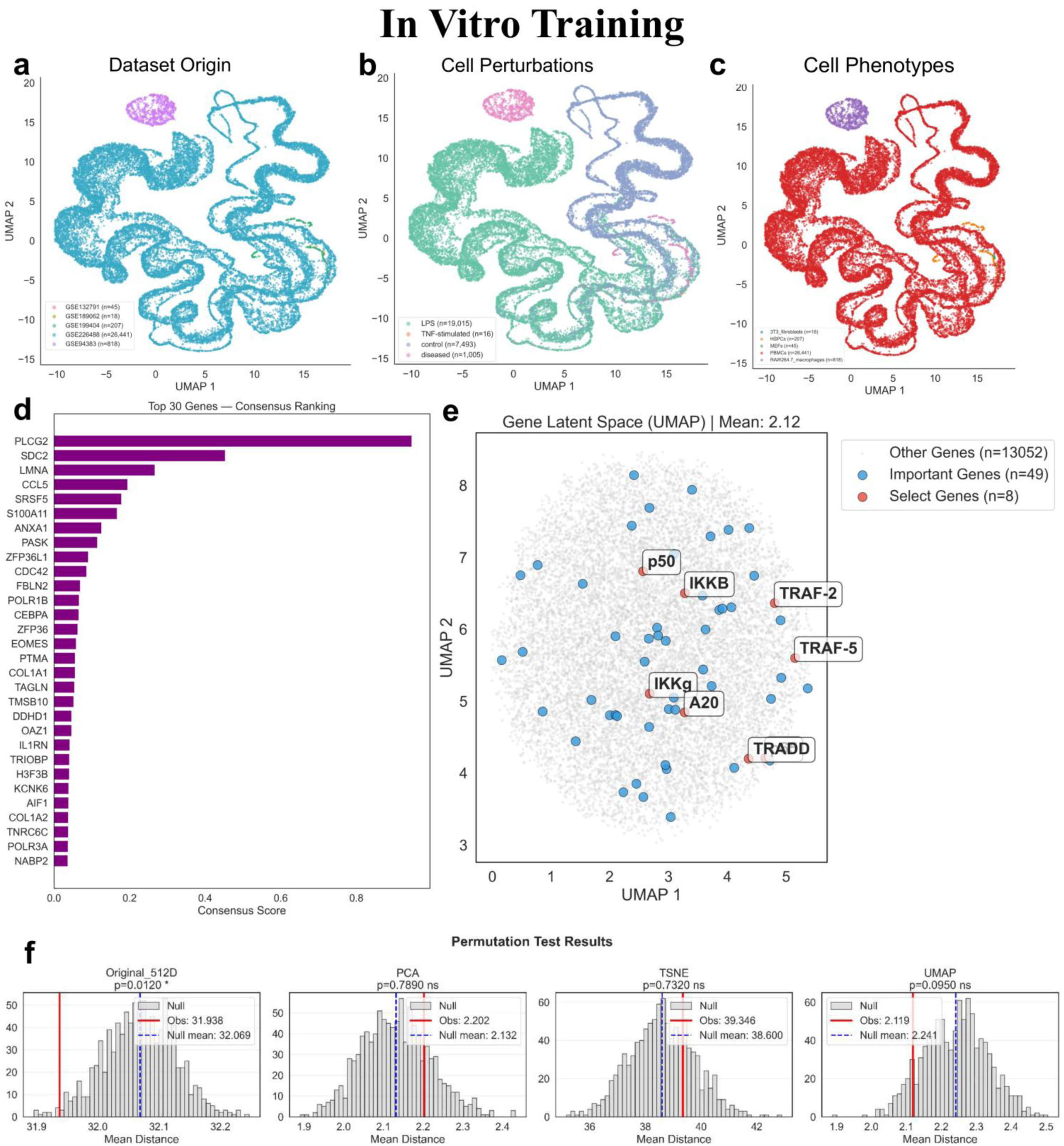
In vitro scGPT foundation model identifies the NFκB gene panel within inflammation-relevant gene latent space. **(a–c)** UMAP visualizations of the fine-tuned scGPT cell embeddings on the curated multi-species in vitro training corpus (27,529 cells post-QC; 31,915 pre-QC), colored by dataset origin (a), perturbation condition (b), and cell phenotype (c). The corpus draws on five GEO accessions: RAW264.7 macrophages (GSE94383; n=818 post-QC), HSPCs/GMPs from GFP-p65/H2B-mCherry mice (GSE199404; n=207 post-QC), 3T3 fibroblast– macrophage co-cultures (GSE189062; n=18 post-QC), MEFs with RelA knockout and complementation (GSE132791; n=45 post-QC), and human PBMCs (GSE226488; n=26,441 post-QC). Perturbation categories are LPS (n=19,015), TNF-stimulated (n=16), control (n=7,493), and diseased (n=1,005), which sum to the 27,529 post-QC cells, and the phenotypes are 3T3 fibroblasts, HSPCs, MEFs, PBMCs, and RAW264.7 macrophages. We fine-tuned the pretrained whole-human scGPT model for binary classification (control vs. inflamed/treated) using a case-insensitive canonical gene vocabulary (∼13,109 genes, after removing mitochondrial, cytoplasmic ribosomal, and canonical hemoglobin artifact families), per-dataset MAD-based QC, mixed multi-species batches, a 70/15/15 stratified train/validation/test split, 20 epochs, learning rate 1×10⁻⁴, effective batch size 32 (via gradient accumulation), and a maximum sequence length of 1,200 genes (Supplementary Fig. 8). **(d)** Bar plot of the top 30 genes identified by the fine-tuned model, ranked by a consensus score computed as the equally weighted mean of three min-max-normalized attribution methods: gradient-based, SHapley Additive exPlanations (SHAP), and perturbation analysis. Leading genes include PLCG2, SDC2, LMNA, CCL5, SRSF5, S100A11, ANXA1, PASK, ZFP36L1, and CDC42. The list is enriched for canonical NFκB-responsive and inflammatory modulators (CCL5, CXCL2, ANXA1, ZFP36L1, ZFP36, CEBPA, IL1RN, AIF1, IFI30) together with ECM and fibroblast-identity genes (COL1A1, COL1A2, FBLN2, TAGLN), consistent with the composition of the corpus. No mitochondrial, ribosomal, or hemoglobin genes appear among the top 30, reflecting the upstream HGNC-curated artifact-gene removal. The full top-50 consensus ranking is provided in Supplementary Table 3. **(e)** Gene latent space visualized as a UMAP projection of the 512-dimensional scGPT gene embeddings, displaying all other genes in grey (n=13,052), the top 50 consensus-ranked reference genes shown in blue (n=49 after one gene lacked a valid embedding coordinate; ranking provided in full in Supplementary Table 3), and the NFκB panel genes in red (n=8; TRAF-5, TRAF-2, TRADD, A20, p50, IKKβ, IKKγ, and p65). The mean UMAP distance between panel genes and the reference-gene neighborhood is 2.12, supporting spatial co-localization within inflammation-relevant feature space. **(f)** Statistical significance of NFκB panel gene proximity to the top-50 consensus-ranked reference gene neighborhood, evaluated by permutation tests in four distance spaces: the native 512-dimensional scGPT embedding (top left), its PCA-reduced projection (top right), and its t-SNE-reduced (bottom left) and UMAP-reduced (bottom right) projections. Each panel compares the observed mean panel-to-reference distance against an empirical null generated by random reference-set resampling. The permutation test reached significance in the native 512-dimensional space (p = 0.0120*), while the PCA (p = 0.7890), t-SNE (p = 0.7320), and UMAP (p = 0.0950) projections did not. Dashed lines indicate p = 0.05 (yellow), p = 0.01 (red), and p = 0.001 (dark red).

Within the learned gene latent space, all 8 NFκB panel genes mapped to the canonical vocabulary and settled near the top 50 consensus-ranked reference genes (the same ranking shown at the top-30 level in Fig. 10d, here extended to 50; Supplementary Table 3), instead of scattering at random among the other 13,052 genes (Fig. 10e; mean UMAP distance 2.12). Formal enrichment testing was mixed. Permutation tests against random reference-gene resampling were significant in the native 512-dimensional scGPT embedding (p = 0.0120) but not in the PCA-reduced (p = 0.7890), t-SNE-reduced (p = 0.7320), or UMAP-reduced (p = 0.0950) projections (Fig. 10f). Carrying the model forward by sequential transfer learning to 15 in vivo lung disease cohorts (Supplementary Fig. 9; Supplementary Table 2) did not recover the NFκB enrichment signal, which we attribute to the far greater transcriptomic heterogeneity of that multi-disease corpus.

The top-50 consensus-ranked list ranges across canonical NFκB-responsive chemokines (CCL5, CXCL2), innate immune regulators (S100A11, ANXA1, AIF1, IFI30, IL1RN, MS4A7), immediate-early AP-1/NFκB-adjacent transcription factors (ZFP36, ZFP36L1, CEBPA, BTG3), ECM and fibrotic-remodeling targets (COL1A1, COL1A2, FBLN2, FN1, TGFBI, BGN, SDC2), cytoskeletal genes (LMNA, TAGLN, ACTB, MYL6, TMSB10, TRIOBP), and signaling components (PLCG2, CDC42, PASK), and it also surfaces the pathway-adjacent regulator IKBIP (IKBKB-interacting protein), which lands in the top 50 even though it is not part of the iseqPLA panel (Supplementary Table 3). Building future iseqPLA probes against these computationally nominated modules (IKBIP in particular, as a functionally NFκB-proximal candidate) is a natural route to foundation-model– guided panel expansion.

## Discussion

We have built a multiplexed 3D spatial interactomics workflow that resolves endogenous NFκB PPIs at single-cell resolution. By coupling iseqPLA to spinning disk confocal microscopy and volumetric reconstruction, it recovers the axial information that conventional 2D PLA discards. Our own 2D-versus-3D comparison bears this out: 3D quantification lowered cell-to-cell variance and sharpened the distinction between nuclear and cytoplasmic signal.

The upstream dissociation kinetics we see in NIH-3T3 fibroblasts line up with the established picture of TNFα-induced Complex I remodeling ^[10]^. Our data add single-cell, subcellular resolution: TNFα collapses the TRAF-5_TRADD and TRAF-5_TRAF-2 populations faster and more completely than IL-1β does, in keeping with the different receptor-proximal architectures of TNFR1 and IL1R ^[1,6]^. The pairwise dynamics that iseqPLA captures are a natural complement to the broad interactome maps produced by proximity labeling ^[13]^.

The effect of ECM coating on NFκB responsiveness has a practical consequence. Matrigel held TNFα-induced activation to 18.2%, well below collagen I (86.8%) and poly-L-lysine (67.2%), most likely through cytokine sequestration, altered integrin signaling, or both; the stronger response on collagen I is what integrin-FAK-IKK crosstalk would predict ^[33]^. Matrigel is often chosen for physiological realism, yet it may quietly dampen cytokine-driven responses and muddy interpretation. The substrate is therefore a variable worth reporting and controlling explicitly in NFκB studies.

Across two independent paradigms, the co-cultures revealed paracrine NFκB amplification. CFASN-conditioned macrophages raised p65 activation in neighboring fibroblasts above the CCL2 controls, in line with the excess IL-1β and TNFα that CF macrophages produce ^[36,39]^. That fibroblasts stay flat to LPS on their own but respond strongly in co-culture shows macrophage-derived paracrine signals are what license NFκB engagement in fibroblasts. Because the same conclusion emerges across independent donor cohorts, two different readouts (phospho-p65 and iseqPLA), and separate laboratories, we are confident that disease-associated conditioning reaches NFκB signaling at the level of supercomplex-scale proximity signals. We note that the cystic fibrosis co-culture observations derive from a single patient donor in each paradigm and should be regarded as preliminary; validation in a multi-donor CF cohort is deferred to future work.

For the panel itself, the scGPT analysis provides exploratory, hypothesis-generating evidence: every NFκB panel gene sits near the top consensus-ranked reference genes in the model’s learned latent space. A permutation test against randomly resampled reference gene sets was significant in the native 512-dimensional embedding (p = 0.0120) but not in the PCA-(p = 0.7890), t-SNE-(p = 0.7320), or UMAP-reduced (p = 0.0950) projections. Our refined 5-dataset in vitro corpus, restricted to cell-line and primary-cell models with defined NFκB perturbations, together with the upstream removal of mitochondrial, ribosomal, and hemoglobin artifact genes, yields a more conservative enrichment estimate than a looser corpus would, the price of trading confounding technical variance for statistical power in a small-panel test. We read the lone significant result (permutation in native space) as suggestive, not confirmatory. Its main value is as a template: a computational framework for nominating candidate PPI panels and modules (ECM remodeling, cytoskeletal reorganization, AP-1/NFκB-adjacent immediate-early regulators, and the pathway-adjacent regulator IKBIP) for experimental validation and future expansion. The loss of significance in the in vivo model (Supplementary Fig. 9) similarly reflects how the heterogeneity of the 15-disease lung corpus dilutes an NFκB-specific signal.

The approach has clear limits. iseqPLA works only in fixed cells; we ran at most four staining cycles; the binary scGPT framing cannot represent continuous activation states; mechanistic dissection of ECM effects and full dose-response experiments were beyond our scope; the co-culture material came from a small number of donors; and the IL-1β dose (1 ng/ml) may fall short of maximal activation in some cell types. All experiments were performed in adherent monolayers and dissociated co-cultures rather than intact tissue, as in our prior tissue-section iseqPLA work [24], because monolayers gave clearer, better-resolved single-cell signal for 3D volumetric quantification. Relative to our previous spatial NFκB study (Zhang et al., APL Bioeng 2024, ref. 42), which mapped NFκB pseudo-signaling at high cell throughput in 2D, the advance here is volumetric: endogenous protein interactions are resolved in 3D at single-cell resolution and interpreted through supercomplex-scale proximity organization. Demonstration of the workflow in intact tissue is deferred to future work. Several constraints temper interpretation: interaction calls rest on a commercial proximity-ligation chemistry without single-probe-omitted, isotype, or genetic specificity controls; the rolling-circle products are better described as co-localized proximity signal than as validated higher-order assemblies and are subject to probe saturation at high interaction density; the phospho-p65 activation metric is a whole-cell co-localization proxy rather than a direct nuclear-translocation readout; only RelA and p105/p50 of the five NFκB subunits were profiled; and error bars are SEM computed across cells from single experiments, so cells are not independent biological replicates. These constraints shape what the study claims. The contribution here is a measurement approach: a way to resolve endogenous multiprotein assemblies inside single cells in three dimensions, together with the descriptors needed to quantify their abundance, size and spatial arrangement. The biological readouts demonstrate what that approach can measure; they are not offered as settled statements about NFκB complex stoichiometry or about cystic fibrosis pathophysiology, and the text throughout has been written to that standard.

Each remaining extension sits outside this study for a specific reason. Genetic ablation of an interaction partner is the standard specificity control, but TRADD, TRAF2, TRAF5, RelA and the IKK subunits are essential nodes of an essential pathway: deleting them compromises viability and rewires the surviving network, so a loss of puncta in the cells that remain cannot be attributed cleanly to probe specificity rather than to an altered cell state or to selection of an escaper population. For the reagents themselves we relied on the validation packages of the commercial proximity-ligation chemistry and of each primary antibody, which provide a vendor-qualified benchmark in the cell types used here; this is a weaker standard than an in-house knockout control and we do not present it as equivalent. A PLA titration defining the linear, unsaturated regime against a single-punctum point-spread reference would sharpen the count-based descriptors further, and the component-area and merged-fraction analysis in Supplementary Fig. 2 is a first step toward it from the existing data. Intact tissue was a deliberate exclusion rather than an oversight: in a tissue section the stimulus reaches different cells at different times through diffusion gradients and several cell types respond in parallel, so a spatiotemporal series cannot be resolved into the per-cell trajectories this method is built to measure. A monolayer and dissociated co-culture, in which every cell receives the same dose at the same moment, is the setting in which the measurement is interpretable; the tissue-section demonstration in our earlier work [24] and the temporal resolution shown here are complementary, and joining them is the natural next step. Biological replicates reported with full distributions and analyzed with a cell-level mixed model, and validation of the CF phenotypes in a multi-donor cohort, require new material and new imaging runs. A representative set of the imaging data is deposited in the BioImage Archive under accession S-BIAD4004.

iseqPLA coupled to 3D confocal imaging offers a spatial interactomics workflow for intracellular signaling supercomplexes. The staggered fixation scheme scales readily to 96-well formats for higher-throughput screening, and extending the approach to patient-derived primary cells, tissue sections, and other signaling pathways should broaden its use for studying immune signaling in health and disease.

## Materials and Methods

### NIH-3T3 mouse fibroblast cell culture

We coated 24-well glass-bottom plates with 0.01% poly-L-lysine and/or collagen I (50 μg/ml, Thermo Fisher A1048301). At 70-80% confluency the cells were seeded onto coverslips, left to adhere overnight, and the next day stimulated with mouse-derived TNFα (10 ng/ml; BioTechne 410-MT), IL-1β (1 ng/ml; BioTechne 401-ML-010), or a DMSO baseline (0.1% v/v). Each well received one condition for 0–105 min at 15-min intervals, inclusive, after which cells were fixed in 4% formaldehyde.

### IMR-90 human fibroblast cell culture

As with the NIH-3T3 cells, we coated 24-well glass-bottom plates with 0.01% poly-L-lysine and/or collagen I (50 μg/ml, Thermo Fisher A1048301); at confluency the cells were seeded onto coverslips, left to adhere overnight, and stimulated the next day with human-derived TNFα (10 ng/ml; BioTechne 10291-TA-050), IL-1β (1 ng/ml; BioTechne 201-LB-005), or a DMSO baseline (0.1% v/v). Each well received one condition for 0–105 min at 15-min intervals, inclusive, after which cells were fixed in 4% formaldehyde for subsequent hybridization chain reaction (HCR) RNA and/or PLA staining.

### Primary CCL2 & CFASN macrophage isolation, transmigration, and co-culture with IMR-90

Dr. Rabindra Tirouvanziam’s laboratory (Department of Pediatrics and Center for CF & Airways Disease Research, Emory University and Children’s Healthcare of Atlanta) generously provided the primary human blood monocytes and their airway-conditioned macrophage derivatives. The two macrophage populations were produced with an established organotypic airway transmigration model developed by the Tirouvanziam group ^[38,39]^, which recapitulates the in vivo recruitment of blood-borne myeloid cells into the lung airway compartment.

In brief, primary human blood monocytes isolated from donor peripheral blood were transmigrated across a well-differentiated human small airway epithelium (H441 cells) that had been grown at the air-liquid interface (ALI) on Alvetex scaffold filters for roughly two weeks to reach full polarization. We inverted the filters so that purified monocytes added to the basal epithelial surface would migrate toward the apical compartment, which held one of two chemoattractants: (1) CCL2 (MCP-1), a canonical monocyte-attracting chemokine that served as the non-disease control, driving chemotaxis without disease-specific airway conditioning; or (2) CF airway supernatant (CFASN), prepared from cystic fibrosis sputum by sequential centrifugation to remove intact cells and bacteria while avoiding denaturing agents such as dithiothreitol (DTT), so as to preserve the native biochemistry of the CF airway. After roughly 10 hours, the cells that had crossed the barrier into the apical compartment were collected as either CCL2-transmigrated macrophages (control, i.e., the monocyte-to-macrophage transition under chemokine guidance alone) or CFASN-transmigrated macrophages (disease-conditioned, i.e., monocytes recruited into the hyperinflammatory CF airway milieu). As Ford et al. previously showed, monocytes transmigrated into CFASN take on a distinct transcriptional and functional phenotype relative to control-transmigrated cells, including altered immune signaling and dysregulated bactericidal activity, mirroring the dysfunctional macrophage phenotype seen in CF airways in vivo ^[39]^.

After transmigration, the CCL2 and CFASN macrophages were moved to the Coskun laboratory and plated with IMR-90 human fibroblasts at a 1:10 macrophage-to-fibroblast ratio on 24-well glass-bottom plates coated with 0.01% poly-L-lysine and collagen I (50 μg/ml, Thermo Fisher A1048301) as above. The co-cultures adhered overnight and reached confluency by day 3, and a separate 24-well plate held IMR-90 cells alone. At day 3, wells were stimulated with 1) LPS 10 ng/ml (InvivoGen tlrl-eblps), 2) TNFα 10 ng/ml (BioTechne 10291-TA-050), 3) IL-1β 1 ng/ml (BioTechne 201-LB-005), or 4) DMSO (0.1% v/v) unstimulated baseline for 0, 30, 60, 120, 240, 480 mins each, and cells were then fixed in 4% formaldehyde for subsequent HCR RNA and/or PLA staining.

### Primary healthy donor & CF-derived monocyte isolation and co-culture with IMR-90

Human subjects were recruited under a protocol approved by the Emory University Institutional Review Board (IRB #2025P010550), and all procedures followed the relevant guidelines and regulations. We obtained written informed consent from every adult participant; for pediatric participants, we obtained written informed assent where appropriate together with parental or guardian consent.

Peripheral blood mononuclear cells (PBMCs) were isolated by Ficoll-Hypaque density-gradient centrifugation from whole blood of people with cystic fibrosis (pwCF) and healthy controls (HC), using protocol modifications optimized for CF samples. From these PBMCs we purified monocytes with the EasySep™ Human Monocyte Isolation Kit (StemCell Technologies, Cat. No. 19359) per the manufacturer’s instructions: PBMCs were incubated with the isolation cocktail and magnetic particles, then negatively selected on a magnetic separator to enrich untouched monocytes. The isolated monocytes were resuspended in CryoStor cryopreservation medium (StemCell Technologies Cat. No. 07930) and stored in liquid nitrogen until use.

Healthy and CF monocytes were moved to the Coskun laboratory, thawed, and plated with IMR-90 human fibroblasts at a 1:10 monocyte-to-fibroblast ratio on 24-well glass-bottom plates coated with 0.01% poly-L-lysine and collagen I (50 μg/ml, Thermo Fisher A1048301) as above. The co-cultures adhered overnight and reached confluency before cytokine stimulation, with a parallel 24-well plate of IMR-90 fibroblast mono-cultures as controls. At confluency, wells were stimulated with 1) LPS 10 ng/ml (InvivoGen tlrl-eblps), 2) TNFα 10 ng/ml (BioTechne 10291-TA-050), 3) IL-1β 1 ng/ml (BioTechne 201-LB-005), or 4) DMSO (0.1% v/v) unstimulated baseline for 0, 30, 60, 120, 240, and 480 minutes each. Cells were then fixed in 4% formaldehyde and processed for 3D iseqPLA staining of A20 & IKKβ and TRAF-5 & TRAF-2 PPIs as described in the Spatial PPI profiling section.

### Spatial PPI profiling in single cells

To detect PPIs, cells were permeabilized in 0.05% Triton X-100 for 10 minutes at room temperature and then processed by iseqPLA ^[24]^ (one PPI per cycle in a single fluorescent channel). Each round proceeded as follows: (1) samples were blocked with Duolink blocking solution to limit nonspecific binding; (2) they were incubated overnight at 4 °C with a pair of target antibodies (one carrying a PLUS oligonucleotide, the other a MINUS oligonucleotide) diluted to the optimal concentration in Duolink antibody diluent; (3) ligase was added at 37 °C for 30 minutes; (4) signal was amplified at 37 °C for 100 minutes; and (5) nuclei were counterstained with DAPI, after which samples were ready for fluorescence imaging. A full iseqPLA cycle (primary antibody incubation (overnight, ∼16 hours), secondary probe binding, ligation, amplification, and washes) takes roughly 20–24 hours; with imaging (∼1–2 hours per 24-well plate), a four-cycle experiment runs about 4–5 days per plate.

To confirm that antibody stripping worked between cycles, we stripped with VectaPlex and reimaged the 647 nm PLA channel; residual signal was negligible (< 5% of pre-stripping intensity) in every experiment, confirming effective removal. To gauge reproducibility, key conditions (e.g., the DMSO control and 45-minute TNFα stimulation) were repeated independently on at least two separate plates processed on different days, and the quantitative metrics (PPI counts per cell, N/C ratios) agreed across replicates, though we note that the staggered single-plate design used for the time courses cannot provide within-experiment biological replication at individual timepoints. Unless stated otherwise, the error bars in all quantitative figures are the standard error of the mean (SEM) across all segmented cells within a given condition and timepoint from a single experiment.

All primary antibodies were diluted in Duolink probe diluent and incubated overnight at 4 °C. The following antibodies and reagents were used: Phospho-S536 p65 (Abcam ab86299, 1:100), p65 (Abcam ab190589, 1:100), p65 (Abcam ab207297, 1:200), p105/p50 (Novus Biologicals NBP2-80872, 1:200), A20 (Novus Biologicals NBP1-77533, 1:100), IKKβ (Abcam ab171364, 1:100), IKKγ/NEMO (Abcam ab230832, 1:100), IκBα (Abcam ab183134, 1:250), TRAF-5 (Novus Biologicals NBP2-80988, 1:250), TRAF-2 (Novus Biologicals NBP1-77248, 1:50), TRADD (Abcam ab238960, 1:100), CD68 (Santa Cruz sc-20060, 1:50), and Phalloidin 555 (Thermo Fisher A34055, 1:400).

### 3D confocal imaging

Imaging used a Cephla Squid spinning disk confocal with a Nikon 60x water lens, following our previously published PRISMS workflow ^[27]^. PLA samples were mounted in 10% glycerol in Wash A buffer, and z-stacks were collected at 0.5 μm spacing over 40 planes and stitched in 3D as a 3×3 grid around each field of view. Each field of view (3×3 stitched grid, 40 z-planes, 3–4 fluorescent channels) took roughly 5–10 minutes to acquire, so a full 24-well plate took about 3-4 days to image.

### Quantitative image analysis and metric definitions

All image analysis ran on custom Python scripts (see Code Availability), with nuclei segmented from DAPI z-stacks using Cellpose ^[43]^. The cytoplasm was taken as the area inside each cell’s segmented boundary but outside the DAPI-defined nucleus. The quantitative metrics used throughout are defined below:

Nuclear-to-cytoplasmic (N/C) p65 ratio (Fig. 2b): the ratio of mean p65 fluorescence intensity within the DAPI-defined nuclear volume to the mean p65 fluorescence intensity within the cytoplasmic volume, computed from 3D volumetric segmentation. Values > 1 indicate nuclear enrichment. For the 2D comparison (Fig. 2b), this ratio was computed from maximum intensity projections of the same z-stacks.

p65 activation score (Pearson r; Figs. 5–6, Supplementary Fig. 5): The Pearson correlation coefficient between Phospho-S536 p65 and total p65 pixel intensities across all pixels within each segmented cell. A Pearson r > 0.5 threshold was used to classify cells as “activated.” This metric captures the spatial co-localization of phosphorylated and total p65 within each cell.

Supercomplex area (μm²; Fig. 4b): The projected area of contiguous above-threshold PLA signal clusters within a maximum intensity projection of the 3D z-stack, measured per cluster and averaged per cell.

Supercomplex count (Fig. 4c): The number of discrete above-threshold PLA signal clusters per cell, identified by intensity thresholding and connected-component analysis.

Sum of dots per cell (Fig. 3b): The total number of individual PLA puncta (RCA products) detected per cell across all z-planes, computed by 3D spot detection.

PPI trajectory analysis (Figs. 1b, 2c): A per-punctum direction-vector field visualizing the spatial relationship between two PLA interaction populations within each cell. For each detected punctum of an upstream interaction (origin; e.g., TRAF-5 & TRAF-2), the k nearest puncta of a downstream interaction (destination; e.g., A20 & IKKβ) were identified in 3D (Z, Y, X) coordinates (k = 5, or fewer when fewer destination puncta were present), their coordinates averaged to a single target point, and the unit direction vector from the origin punctum to that target computed (scikit-learn NearestNeighbors). The resulting normalized vectors are rendered as white arrows overlaid on the 3D reconstruction. This metric is a spatial-directionality visualization between interaction populations and does not represent temporal tracking.

### Statistics

Unless stated otherwise, error bars represent the standard error of the mean (SEM) computed across all segmented cells of a single experiment, and n denotes the number of cells, reported per panel. Pairwise per-cell comparisons against the matched DMSO control use two-sided Mann–Whitney U tests, with a linear mixed-effects model (fixed effect: stimulation; random intercept: field of view) as a sensitivity analysis, and p-values are adjusted by Benjamini–Hochberg correction across the comparisons within each figure panel. The scGPT analyses use the permutation, Kolmogorov–Smirnov, and Mann–Whitney tests indicated in the corresponding panels. Cell numbers are reported at two levels: the number of segmented cells imaged in an experiment, given in the cell-count heatmaps (Supplementary Figs. 1 and 7), and the number of cells entering each quantification panel, given in that panel legend; the two differ where a cell carried no measurable signal for the marker in question. All experiments are single biological replicates and individual cells are not independent measurements, which we state as a limitation (Discussion).

### scGPT foundation model training

The scGPT (single-cell Generative Pretrained Transformer) foundation model ^[40]^ was trained in a two-stage transfer-learning pipeline. For the in vitro model, we first assembled a multi-species training corpus, intersecting the gene vocabularies of four curated mouse in vitro NFκB datasets and one human PBMC dataset (Supplementary Table 1) by case-insensitive symbol matching; an alternative MGI HOM_MouseHumanSequence ortholog-based intersection gave equivalent results. Three technical-artifact gene families (mitochondrial transcripts (MT-prefix), cytoplasmic ribosomal proteins (HGNC group 1054, 90-gene allowlist), and canonical hemoglobin subunits (HGNC group 940, 10-gene allowlist)) were removed before training through explicit curated allowlists rather than prefix-based regex, which avoids false positives such as HBEGF, HBO1, RPS6KA1–6, and RPS6KB1–2, leaving a final canonical vocabulary of about 13,109 UPPER-case gene symbols. Quality control followed the recipe at sc-best-practices.org: cells were filtered by 5-MAD thresholds on log1p total counts, log1p gene count, and percent-top-20 expression; by a 3-MAD threshold plus a 20% absolute ceiling on percent mitochondrial reads; and by a 50-gene-per-cell absolute floor. Outlier calling by MAD was done per dataset, not globally, so the majority human dataset would not dominate the median/MAD estimates, and groups of fewer than 10 cells skipped MAD filtering entirely. We then fine-tuned the pretrained whole-human scGPT checkpoint for binary classification (control vs. inflamed/treated) on the QC-filtered corpus, using a learning rate of 1×10⁻⁴, an effective batch size of 32 (batch size 8 with 4-step gradient accumulation), a maximum input sequence length of 1,200 genes, a 70/15/15 stratified train/validation/test split, 20 epochs with early stopping (patience 5), weight decay 0.01, 500 warmup steps, and gradient clipping at 1.0 (Supplementary Fig. 8).

For the in vivo model, we further fine-tuned the in vitro checkpoint on the curated in vivo lung-disease corpus (Supplementary Table 2) with multi-GPU DataParallel training ^[44]^. DataParallel splits each batch across the available GPUs, replicates the model on every device, and synchronizes gradients after each forward-backward pass, which makes training feasible on datasets too large for one GPU. The hyperparameters were a learning rate of 5×10⁻⁵, batch size 16 (effective per-GPU batch size 8), maximum sequence length 3,000 genes, 10 epochs, balanced class subsampling, and best-model selection by F1 score.

Gene-level analysis of the fine-tuned scGPT model comprised a single consensus attribution pipeline serving two purposes: identifying the genes most informative for the model’s binary classification decisions, and defining a reference gene set against which to test enrichment of the iseqPLA panel in the learned gene latent space. Three complementary post hoc attribution methods were applied to the trained classifier: (1) gradient-based attribution, which computes the gradient of the classification loss with respect to the gene embedding inputs; (2) SHapley Additive exPlanations (SHAP) ^[45]^, which estimates each gene’s marginal contribution to the prediction via Shapley value approximation; and (3) perturbation analysis, which measures the change in predicted probability when individual gene inputs are masked. Each method’s per-gene score was min-max normalized to its corpus-wide maximum, and the three normalized scores were averaged with equal weights to produce a single consensus score per gene. The top 30 consensus-ranked genes are displayed in Fig. 10d, and the full top 50 list is provided in Supplementary Table 3. The 50 highest-ranked genes also constituted the “important gene” reference neighborhood against which the NFκB panel genes were compared in the gene latent space (Fig. 10e); one reference gene lacked a valid embedding coordinate and is therefore shown as n = 49 in panel e. We tested whether the pairwise distances from each NFκB panel gene to the 50 reference genes, computed in four distance spaces (the 512-dimensional scGPT embedding and its PCA, TSNE, and UMAP projections) were smaller than expected by chance using three complementary statistical tests implemented in SciPy: a Mann-Whitney U test and a two-sample Kolmogorov-Smirnov test comparing the panel-to-reference distance distribution against the panel-to-all-other-genes distribution, and a permutation test in which the reference gene set was randomly resampled to construct an empirical null distribution (Fig. 10f; Supplementary Fig. 9f). Gene-name harmonization between the NFκB panel (TRAF-5, TRAF-2, TRADD, A20, p105/p50, IKKβ, IKKγ, p65) and the corpus vocabulary was performed using a curated alias dictionary (e.g., p65→RELA/Rela; A20→TNFAIP3/Tnfaip3; IKKβ→IKBKB/Ikbkb; IKKγ→IKBKG/Ikbkg/NEMO; p105/p50→NFKB1/Nfkb1; TRAF-5→TRAF5/Traf5; TRAF-2→TRAF2/Traf2; TRADD→TRADD/Tradd) with case-insensitive exact matching and length-gated partial matching as fallbacks.

## Code Availability

The code used to generate the figures shown in this study can be found here: https://github.com/coskunlab/3D-iseqPLA

## Data Availability Statement

A representative set of registered, stitched images from the CCL2 macrophage and IMR-90 fibroblast co-culture experiment of Fig. 5 (plate 006A, LPS-and vehicle-treated wells at 60 min; cycle 1 immunofluorescence channels for DAPI, phospho-S536 p65 and total p65, and cycle 4 proximity-ligation channels for DAPI and A20 & IKKγ) is deposited in the BioImage Archive under accession S-BIAD4004 (https://www.ebi.ac.uk/biostudies/bioimages/studies/S-BIAD4004). The complete image set runs to several terabytes of stitched 3D volumes and is impractical to deposit in full, so the remaining images and the single-cell measurement tables can be requested from the corresponding author. Additional sample data are available at https://doi.org/10.6084/m9.figshare.31385329

## Supporting Information

Supporting Information is available with the manuscript.

## Author Contributions

A.F.C and N.Z. conceived the study. N.Z performed experiments, analyzed data, developed computational pipelines, and wrote the manuscript. C.L.-T. and H.O. provided primary human monocytes and CF patient-derived cells, performed transmigration experiments, and contributed to manuscript revision. S.S., D.N., and L.R. performed experiments. R.T. supervised the transmigration model and macrophage generation and provided resources, including funding. B.K. supervised monocyte isolation and provided clinical samples and resources, including funding. A.F.C. conceived and supervised the study, secured funding, and edited the manuscript. All authors reviewed and approved the final manuscript.

## Acknowledgments

A.F.C. holds a Career Award at the Scientific Interface from the Burroughs Wellcome Fund and Bernie-Marcus Early-Career Professorship. A.F.C. was supported by start-up funds from the Georgia Institute of Technology and Emory University. Research reported in this study was supported by the National Institutes of Health under award numbers R35GM151028, R33CA291197, and T32GM142616. This work is partially supported by the National Science Foundation CAREER under grant number 2338935 and by H.O.’s Cystic Fibrosis Foundation Postdoctoral Fellowship.

