## Supplemental Information for "3D Spatial Interactomics Maps the Dynamics of NF-κB Multiprotein Signalosomes in Single Cells"

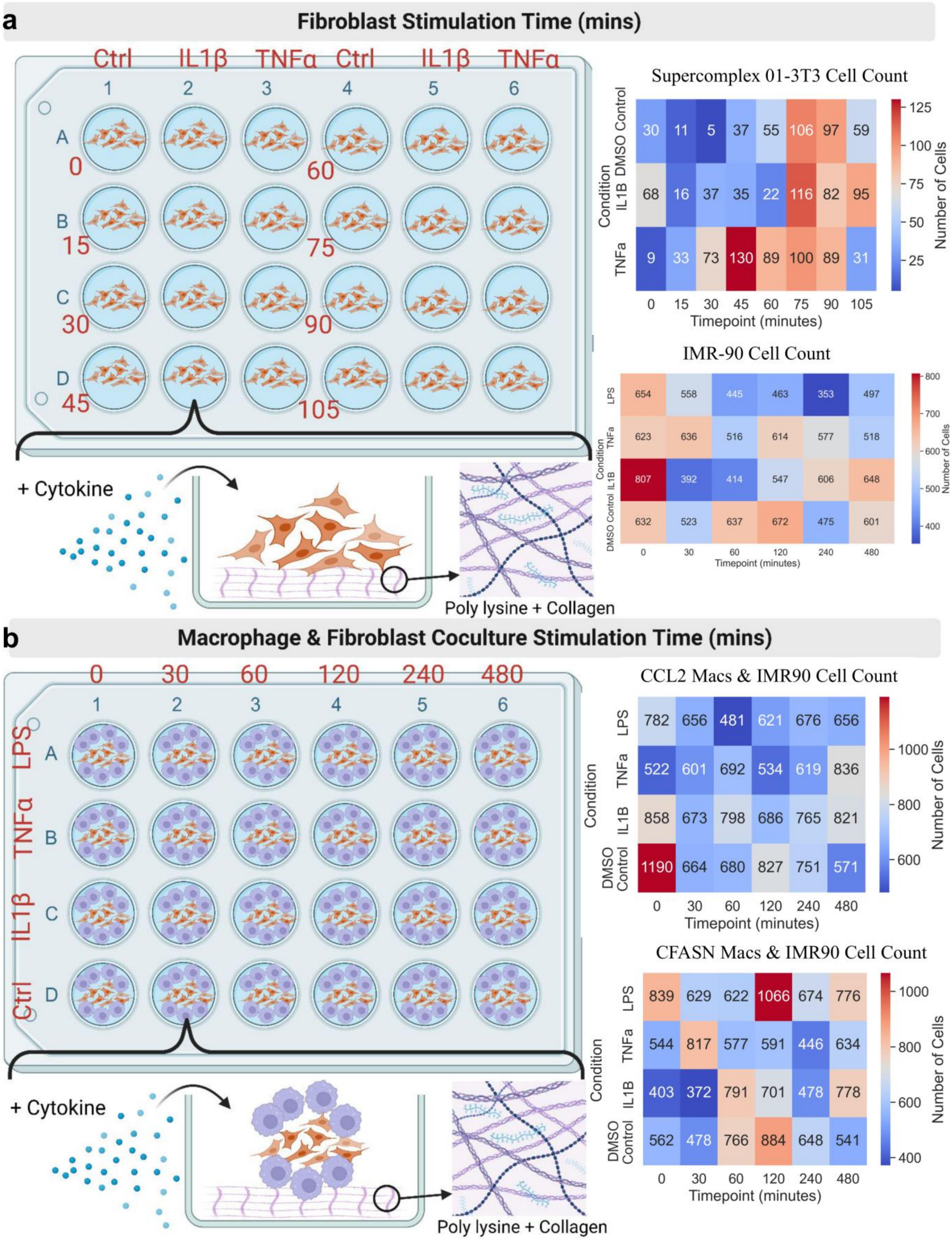
a
Fibroblast Stimulation Time (mins)
Ctrl IL1β TNFα Ctrl IL1β TNFα
1 2 3 4 5 6
A B C D
0 15 30 45 60 75 90 105
+ Cytokine
Poly lysine + Collagen
Supercomplex 01-3T3 Cell Count
DMSO Control: 30 11 5 37 55 106 97 59
IL1B: 68 16 37 35 22 116 82 95
TNFa: 9 33 73 130 89 100 89 31
Condition
Timepoint (minutes)
0 15 30 45 60 75 90 105
Number of Cells
125 100 75 50 25
IMR-90 Cell Count
LPS: 654 558 445 463 353 497
TNFa: 623 636 516 614 577 518
IL1B: 807 392 414 547 606 648
DMSO Control: 632 523 637 672 475 601
Condition
Timepoint (minutes)
0 30 60 120 240 480
Number of Cells
800 700 600 500 400
b
Macrophage & Fibroblast Coculture Stimulation Time (mins)
0 30 60 120 240 480
1 2 3 4 5 6
LPS A
TNFα B
IL1β C
Ctrl D
+ Cytokine
Poly lysine + Collagen
CCL2 Macs & IMR90 Cell Count
LPS: 782 656 481 621 676 656
TNFa: 522 601 692 534 619 836
IL1B: 858 673 798 686 765 821
DMSO Control: 1190 664 680 827 751 571
Condition
Timepoint (minutes)
0 30 60 120 240 480
Number of Cells
1000 800 600
CFASN Macs & IMR90 Cell Count
LPS: 839 629 622 1066 674 776
TNFa: 544 817 577 591 446 634
IL1B: 403 372 791 701 478 778
DMSO Control: 562 478 766 884 648 541
Condition
Timepoint (minutes)
0 30 60 120 240 480
Number of Cells
1000 800 600 400

Supplementary Fig. 1. Experimental design for cytokine stimulation time courses in fibroblast mono-cultures and macrophage-fibroblast co-cultures.

Schematic of 24-well plate layouts and stimulation conditions used across experiments.

**(a)** Fibroblast-only stimulation design. NIH-3T3 fibroblasts seeded on poly-L-lysine and collagen I-coated glass-bottom plates were stimulated with IL1β (1 ng/ml), TNFα (10 ng/ml), or a DMSO vehicle control at staggered time points (0–105 min in 15-min intervals) and then fixed together for downstream PLA and imaging. The right inset shows cytokine being added to adherent fibroblasts on a poly-L-lysine/collagen I ECM substrate. The cell-count heatmaps (right) give the number of segmented cells per well for two parallel mono-culture experiments: NIH-3T3 mouse fibroblasts (top), used for iseqPLA supercomplex profiling, and IMR-90 human fibroblasts (bottom), used for phospho-p65 imaging. This provides a quality-control check that sampling was adequate across conditions. Created with BioRender.com.

**(b)** Macrophage-fibroblast co-culture stimulation design. Two macrophage populations, CCL2-transmigrated control macrophages (chemokine-only, non-disease) and CFASN-transmigrated, CF airway-conditioned macrophages, were each combined with IMR-90 fibroblasts at a 1:10 ratio on poly-L-lysine and collagen I-coated plates. The co-cultures were stimulated with LPS (10 ng/ml), TNFα (10 ng/ml), IL1β (1 ng/ml), or a DMSO control for 0, 30, 60, 120, 240, or 480 min and then fixed. The right inset shows cytokine being added to the mixed macrophage (round, blue) and fibroblast (elongated, orange) co-culture on a collagen-coated substrate. As above, the cell-count heatmaps (right) give segmented cells per well as a quality-control check on sampling across conditions. Created with BioRender.com.

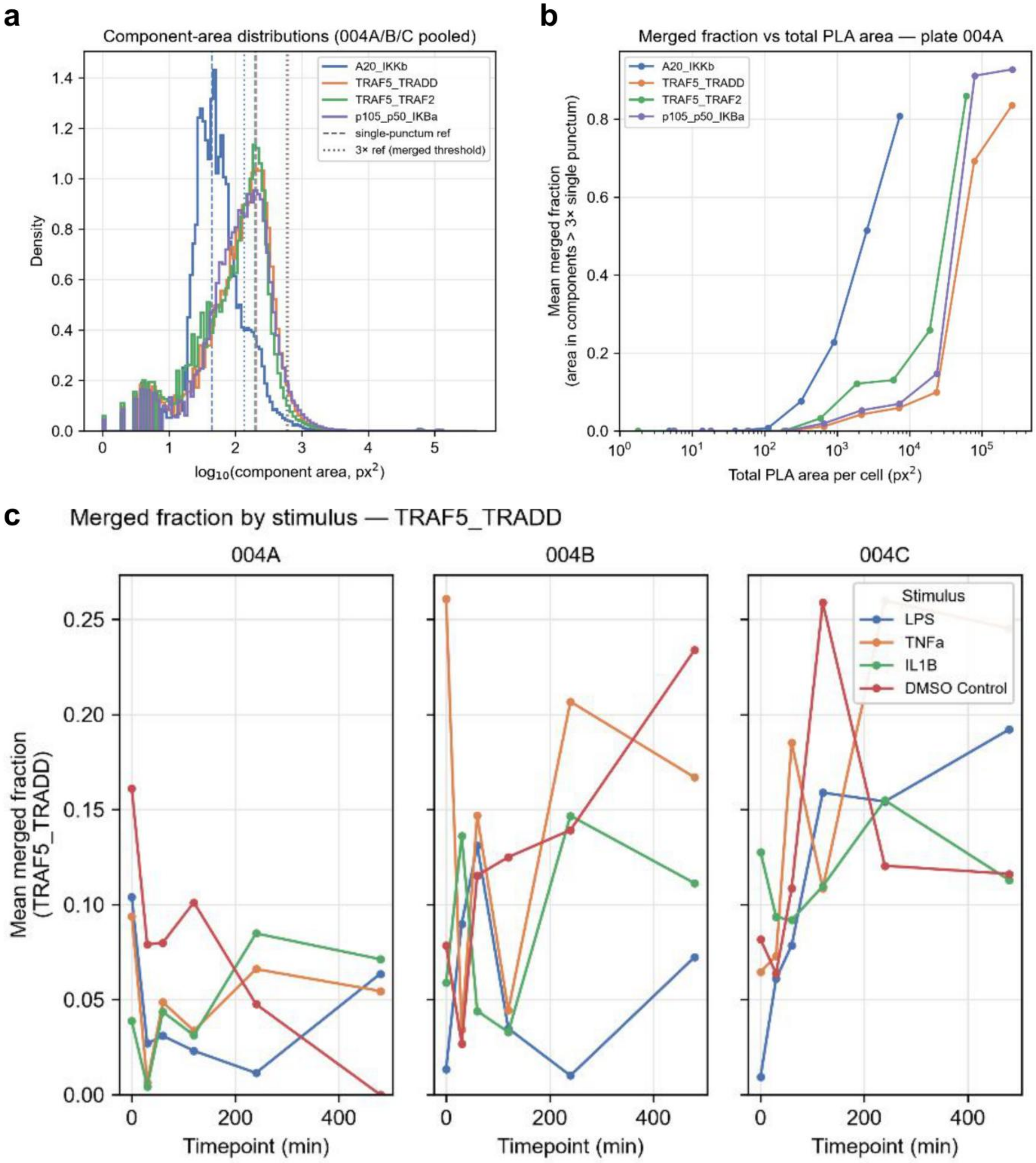
a
Component-area distributions (004A/B/C pooled)
A20_IKKb
TRAF5_TRADD
TRAF5_TRAF2
p105_p50_IKBa
single-punctum ref
3× ref (merged threshold)
Density
log10(component area, px2)
b
Merged fraction vs total PLA area — plate 004A
A20_IKKb
TRAF5_TRADD
TRAF5_TRAF2
p105_p50_IKBa
Mean merged fraction
(area in components > 3× single punctum)
Total PLA area per cell (px2)
c
Merged fraction by stimulus — TRAF5_TRADD
004A
004B
004C
Stimulus
LPS
TNFa
IL1B
DMSO Control
Mean merged fraction
(TRAF5_TRADD)
Timepoint (min)

Supplementary Fig. 2. Component-area distributions and saturation-regime analysis of PLA signal in monocyte and IMR-90 fibroblast co-cultures.

Rolling circle amplification (RCA) products measure roughly 0.5–1 μm across, so neighboring products merge at high interaction density and connected-component counts underestimate the true number of interactions. The linear range was characterized from the component-area distributions of plates 004A, 004B, and 004C (1,642,699 components in total).

(a) Distribution of connected-component areas for each probe pair, pooled across the three plates. The single-punctum reference area for each pair (dashed) is the dominant mode of the $\log_{10}$ area distribution: 45 $px^2$ (0.559 $\mu m^2$) for A20_IKKβ, 207 $px^2$ (2.596 $\mu m^2$) for TRAF-5_TRADD, 194 $px^2$ (2.439 $\mu m^2$) for TRAF-5_TRAF-2, and 201 $px^2$ (2.527 $\mu m^2$) for p105_p50_IκBα. Components exceeding 3× the reference (dotted) were scored as merged. The smallest peak in each distribution (2–6 px) is threshold debris present even in DMSO and 0 min controls, and was not used as the reference.

(b) Mean per-cell merged fraction, the proportion of a cell's total PLA area contained in merged components, against total PLA area per cell in logarithmically spaced bins for plate 004A. Bins with fewer than 5 cells are omitted. The merged fraction rises from ~0% in the lowest-signal cells to 81–93% in the densest cells on plate 004A, and to 81–98% across plates 004A–C. Component counting is therefore approximately linear only at low signal and progressively undercounts as density increases.

(c) Mean merged fraction for TRAF-5_TRADD across stimulation conditions and timepoints, shown separately for each plate. Stimulation increases the number of components per cell rather than their size, so the merged fraction is comparable across conditions and the comparisons in Figs. 7–9 are not driven by differential saturation.

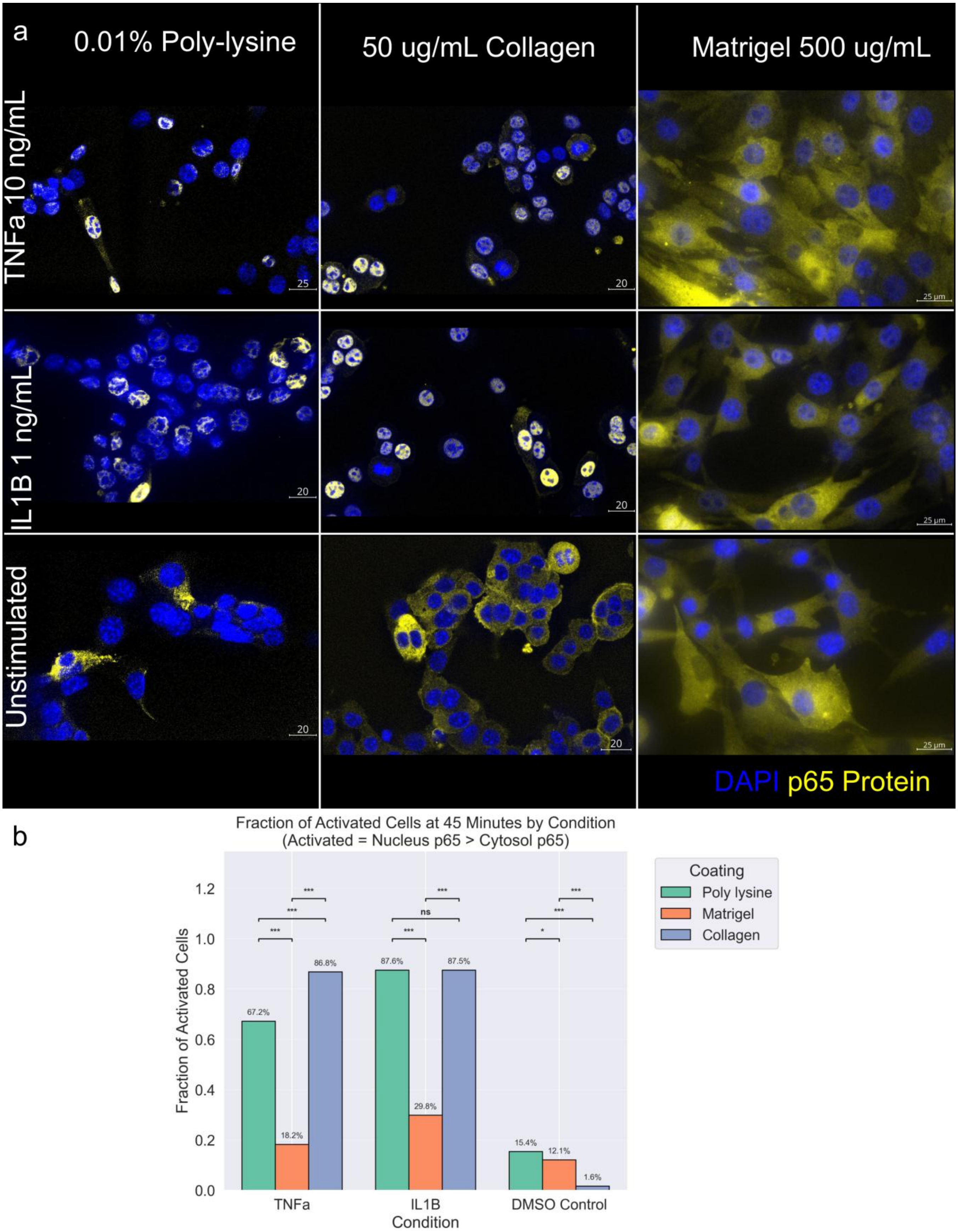
a
0.01% Poly-lysine
50 ug/mL Collagen
Matrigel 500 ug/mL
TNFa 10 ng/mL
IL1B 1 ng/mL
Unstimulated
25
20
25 µm
DAPI p65 Protein
b
Fraction of Activated Cells at 45 Minutes by Condition
(Activated = Nucleus p65 > Cytosol p65)
Fraction of Activated Cells
Coating
Poly lysine
Matrigel
Collagen
67.2%
18.2%
86.8%
87.6%
29.8%
87.5%
15.4%
12.1%
1.6%
***
ns
*
TNFa
IL1B
DMSO Control
Condition

Supplementary Fig. 3. Extracellular matrix coating dictates cytokine accessibility and NF-κB p65 nuclear activation in NIH-3T3 mouse fibroblasts.

**(a)** Representative maximum-intensity-projection confocal images of NIH-3T3 fibroblasts 45 minutes after stimulation with TNFα (10 ng/ml, top row), IL-1β (1 ng/ml, middle row), or a DMSO unstimulated control (bottom row), shown across three substrate coatings: 0.01% poly-L-lysine (left column), 50 μg/ml collagen I (center column), and 500 μg/ml Matrigel (right column). Each two-channel image shows DAPI (blue, nuclei) and p65 protein (yellow). Poly-L-lysine wells adhered fewer cells and, under stimulation, showed weaker cytoplasmic p65 signal, whereas collagen I and Matrigel supported denser growth with distinct p65 subcellular patterns. Matrigel wells retained elevated cytoplasmic p65 even under cytokine stimulation, in line with ECM-mediated control of cytokine access and receptor engagement. Scale bars: 20–25 μm as indicated per panel.

**(b)** Bar graph of the fraction of p65-activated cells (nuclear p65 > cytosolic p65) at 45 minutes, shown for TNFα, IL-1β, and DMSO control on each coating. Under TNFα, collagen I (blue) and poly-L-lysine (green) activated significantly larger fractions of cells (86.8% and 67.2%, respectively) than Matrigel (18.2%), which sharply blunted cytokine-induced p65 nuclear translocation (*** $p < 0.001$, * $p < 0.05$; ns = not significant). Under IL-1β, collagen I and poly-L-lysine reached comparably high fractions (~87%), while DMSO baselines stayed low on every coating (1.6–15.4%), with poly-L-lysine showing slightly more spontaneous activation. The takeaway is that ECM composition strongly governs NF-κB pathway accessibility and should be controlled carefully in inflammatory-signaling studies.

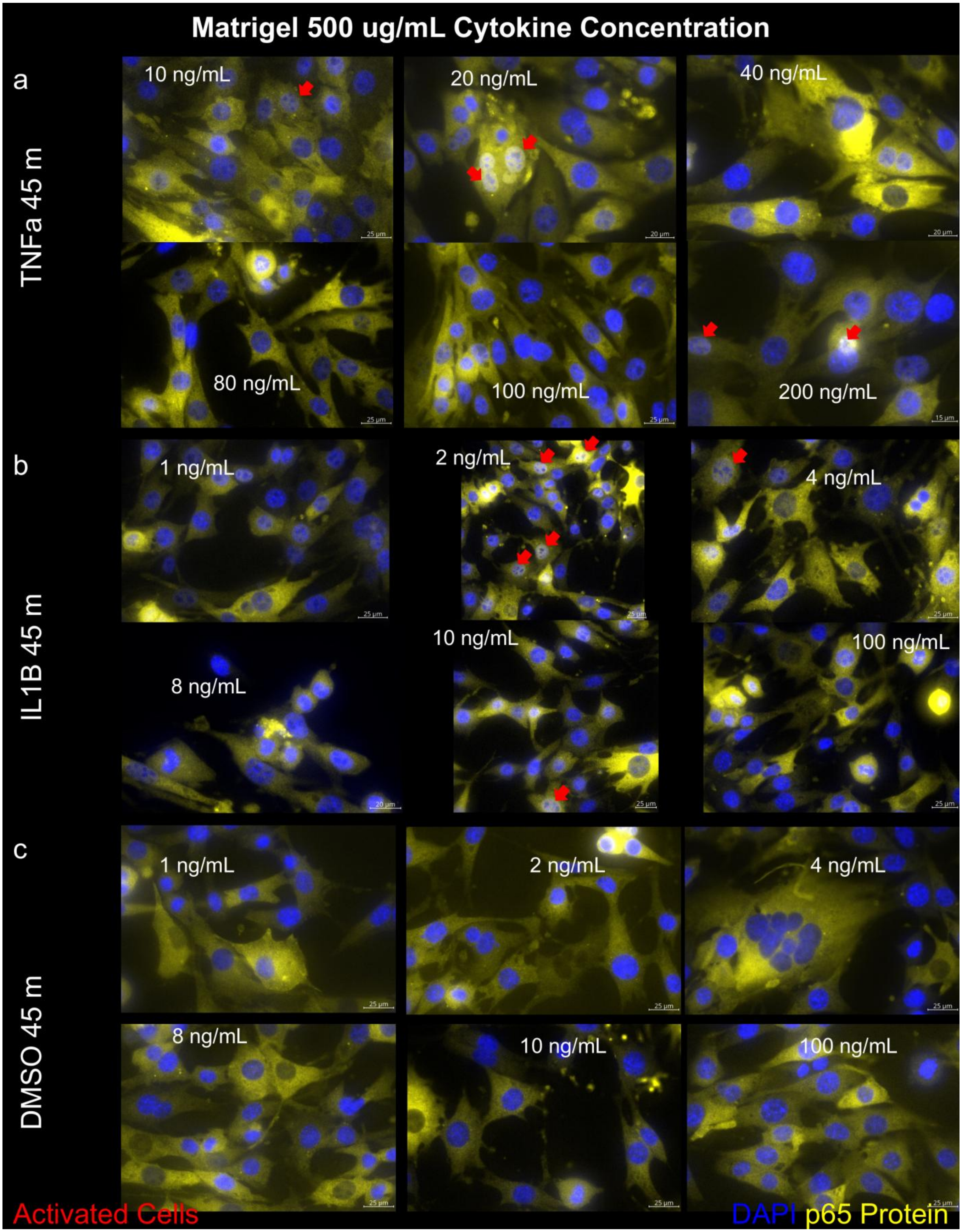
Matrigel 500 ug/mL Cytokine Concentration
a
TNFa 45 m
10 ng/mL
20 ng/mL
40 ng/mL
80 ng/mL
100 ng/mL
200 ng/mL
b
IL1B 45 m
1 ng/mL
2 ng/mL
4 ng/mL
8 ng/mL
10 ng/mL
100 ng/mL
c
DMSO 45 m
1 ng/mL
2 ng/mL
4 ng/mL
8 ng/mL
10 ng/mL
100 ng/mL
Activated Cells
DAPI p65 Protein

Supplementary Fig. 4. Raising the cytokine concentration does not overcome Matrigel-mediated suppression of NF-κB p65 nuclear translocation in NIH-3T3 fibroblasts.

**(a)** Representative maximum-intensity-projection confocal images of NIH-3T3 fibroblasts grown on a 500 μg/ml Matrigel substrate and fixed 45 minutes after stimulation with a TNFα concentration series (10, 20, 40, 80, 100, and 200 ng/ml). DAPI marks nuclei (blue) and total p65 is in yellow; red arrowheads mark activated cells (nuclear p65 exceeding cytosolic p65). Nuclear p65 enrichment stays modest across the entire TNFα range, and most p65 remains cytoplasmic even at 200 ng/ml.

**(b)** As in (a), but for IL-1β across 1, 2, 4, 8, 10, and 100 ng/ml. A modest activated fraction appears at the lower concentrations and does not grow substantially at higher doses.

**(c) DMSO vehicle controls at volumes matched to the (a) and (b) series; each panel is labelled with the concentration of the (a) or (b) condition whose vehicle volume it matches, not with a DMSO concentration.** Activated cells are rare in every condition. Scale bars, 25 μm. Read together with Supplementary Fig. 3, these images suggest that Matrigel caps cytokine-driven p65 activation in a way that extra ligand does not readily overcome, pointing to reduced cytokine access or receptor engagement within the dense matrix rather than a simple rightward shift in sensitivity.

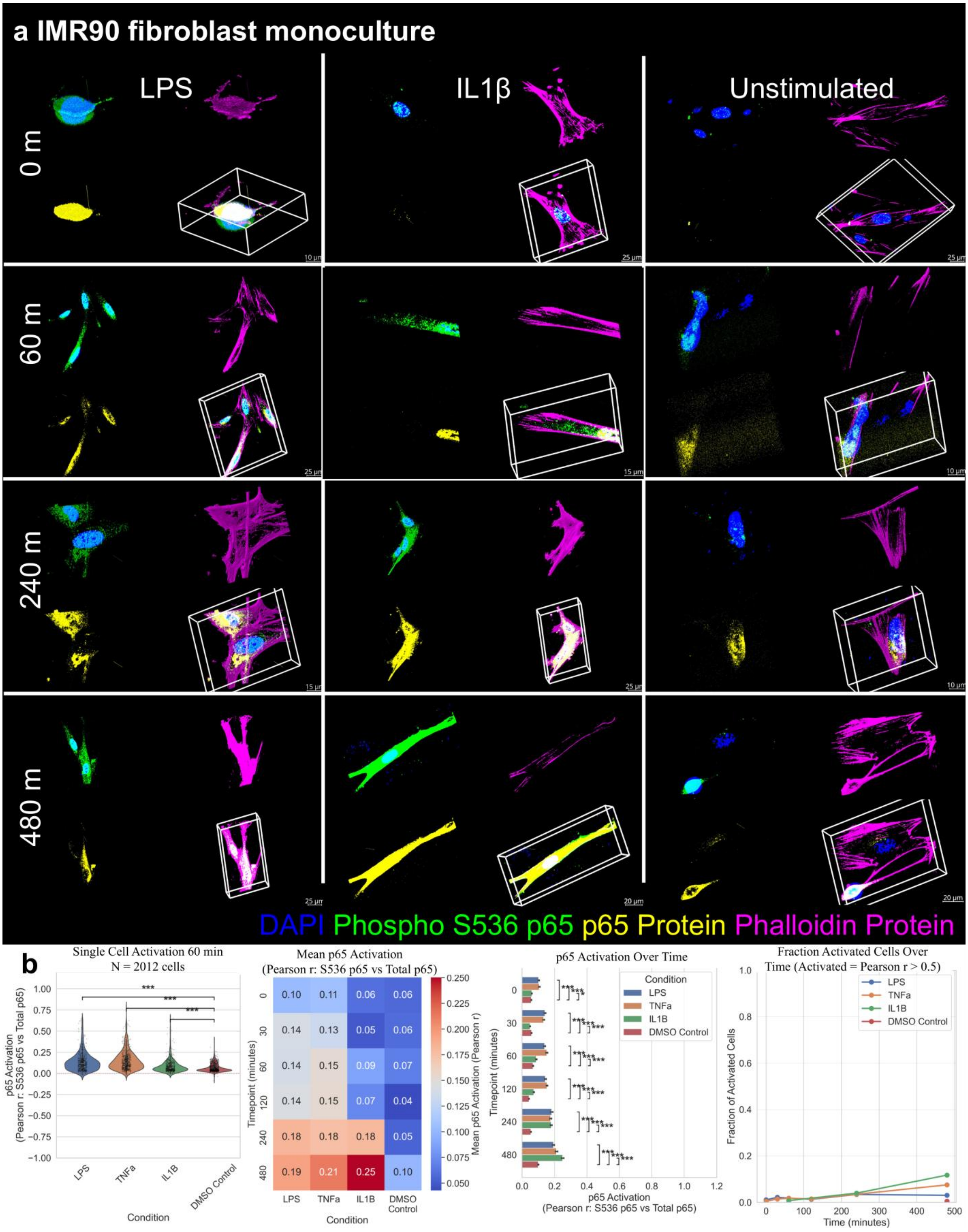
a IMR90 fibroblast monoculture
LPS
IL1β
Unstimulated
0 m
60 m
240 m
480 m
DAPI Phospho S536 p65 p65 Protein Phalloidin Protein
b
Single Cell Activation 60 min
N = 2012 cells
p65 Activation (Pearson r: S536 p65 vs Total p65)
Condition
LPS
TNFa
IL1B
DMSO Control
Mean p65 Activation
(Pearson r: S536 p65 vs Total p65)
Timepoint (minutes)
Mean p65 Activation (Pearson r)
p65 Activation Over Time
p65 Activation
(Pearson r: S536 p65 vs Total p65)
Fraction Activated Cells Over
Time (Activated = Pearson r > 0.5)
Fraction of Activated Cells
Time (minutes)

Supplementary Fig. 5. NFκB p65 activation dynamics in corresponding IMR-90 fibroblast mono-cultures.

**(a)** Representative 3D rendered images of IMR-90 fibroblast mono-cultures (no macrophages) at 0, 60, 240, and 480 min post-stimulation with LPS (10 ng/ml, left column), IL1β (1 ng/ml, center column), or DMSO unstimulated control (right column). In total, 13,408 cells were imaged across all timepoints and conditions, with DAPI (blue), Phospho-S536 p65 (green), total p65 protein (yellow), and Phalloidin (magenta); the elongated fibroblast morphology is well resolved in 3D at every timepoint. Relative to the CCL2 (Fig. 5) and CFASN (Fig. 6) macrophage co-cultures, these mono-cultures show weaker, more transient p65 activation, especially under LPS, as expected given the LPS insensitivity of fibroblasts without paracrine macrophage signals, and IL1β elicits the strongest phospho-p65 response in mono-culture. Scale bars as indicated per panel.

**(b)** Quantitative single-cell analyses of p65 activation in IMR-90 fibroblast mono-cultures. The violin panel shows the 60-minute subset (n = 2,012 cells: LPS 445, TNFα 516, IL1β 414, DMSO 637); the heatmap and time courses draw on all 13,408 segmented cells across the six timepoints, with per-condition counts given in Supplementary Fig. 1a. Far left: violin plots of p65 activation scores (Pearson r of Phospho-S536 p65 vs. total p65) at 60 minutes across LPS, TNFα, IL1β, and DMSO control, with significance annotations; all three stimulations differed significantly from the DMSO control at 60 minutes (*** $p < 0.001$, two-sided Mann–Whitney U with Benjamini–Hochberg adjustment), and IL1β also exceeded LPS; the absolute differences are nonetheless small relative to the cell-to-cell spread, and every condition stayed in the low activation range. Center left: heatmap of mean p65 activation across all four conditions (rows) and timepoints 0–480 min (columns); IL1β climbed in the late phase (0.18 at 240 min, 0.25 at 480 min), while LPS (0.10–0.19), TNFα (0.11–0.21), and DMSO control (0.04–0.10) stayed uniformly low. Center right: mean p65 activation over time for all four conditions, with error bars and pairwise comparisons, confirming that IL1β diverges from the rest at later timepoints. Far right: fraction of p65-activated cells (Pearson $r > 0.5$) over time; every condition, IL1β included, held activated fractions near zero across the 0–480 minute window, a sharp contrast to the macrophage co-cultures (Figs. 5, 6).

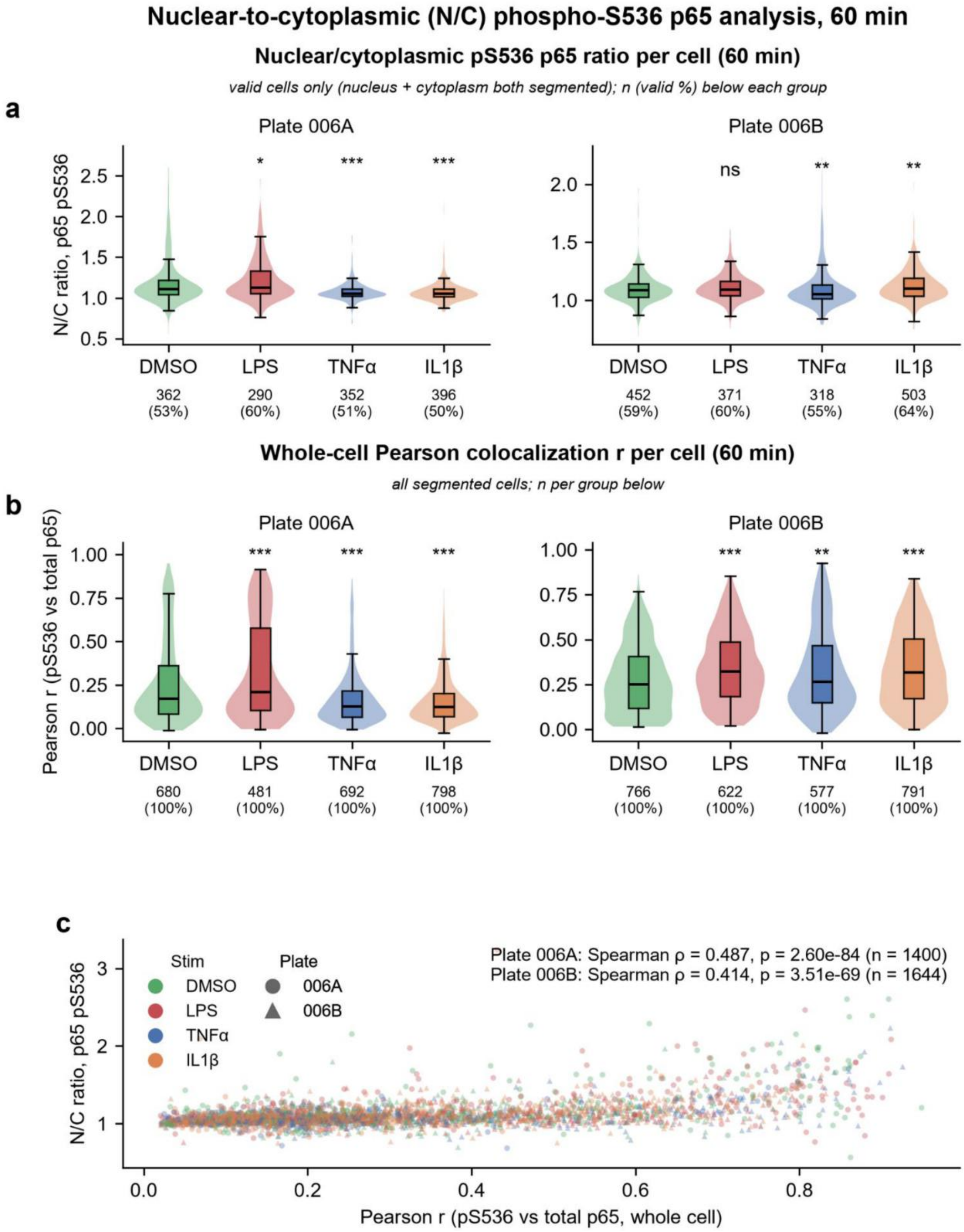

Nuclear-to-cytoplasmic (N/C) phospho-S536 p65 analysis, 60 min
Nuclear/cytoplasmic pS536 p65 ratio per cell (60 min)
valid cells only (nucleus + cytoplasm both segmented); n (valid %) below each group
a
Plate 006A
N/C ratio, p65 pS536
DMSO 362 (53%)
LPS 290 (60%) *
TNFα 352 (51%) ***
IL1β 396 (50%) ***
Plate 006B
DMSO 452 (59%)
LPS 371 (60%) ns
TNFα 318 (55%) **
IL1β 503 (64%) **
Whole-cell Pearson colocalization r per cell (60 min)
all segmented cells; n per group below
b
Plate 006A
Pearson r (pS536 vs total p65)
DMSO 680 (100%)
LPS 481 (100%) ***
TNFα 692 (100%) ***
IL1β 798 (100%) ***
Plate 006B
DMSO 766 (100%)
LPS 622 (100%) ***
TNFα 577 (100%) **
IL1β 791 (100%) ***
c
Stim
DMSO
LPS
TNFα
IL1β
Plate
006A
006B
Plate 006A: Spearman ρ = 0.487, p = 2.60e-84 (n = 1400)
Plate 006B: Spearman ρ = 0.414, p = 3.51e-69 (n = 1644)
N/C ratio, p65 pS536
Pearson r (pS536 vs total p65, whole cell)

Supplementary Fig. 6. Nuclear-to-cytoplasmic phospho-S536 p65 ratios at 60 minutes in macrophage and IMR-90 fibroblast co-cultures.

Single-cell quantification of phospho-S536 p65 (pS536) nuclear translocation at 60 min in the co-cultures of Figs. 5 and 6 (plates 006A and 006B), comparing a direct nuclear-to-cytoplasmic (N/C) intensity ratio with the whole-cell Pearson co-localization coefficient used in the main text.

(a) Per-cell N/C ratio of mean pS536 intensity (nuclear mean divided by cytosolic mean) for DMSO, LPS, TNFα, and IL-1β. A valid ratio requires both a nuclear and a cytosolic compartment in the segmentation, which holds for 52.8% of cells on plate 006A (1,400 of 2,651) and 59.7% on plate 006B (1,644 of 2,756); the remainder are cytosol-only segmentations and are excluded here. Valid n and the corresponding percentage are printed below each group. Violins show the kernel density and boxes the median, interquartile range, and 1.5× IQR whiskers, with outliers hidden. The N/C ratio does not rise with stimulation, most likely because IKK activation drives de novo pS536 phosphorylation in the cytosol rather than because translocation fails.

(b) The same cells, without the segmentation requirement, scored by whole-cell Pearson correlation between pS536 and total p65 intensity. This metric separates LPS from DMSO on both plates (Benjamini–Hochberg-adjusted Mann–Whitney U, $p = 4.1 \times 10^{-4}$ for 006A and $p = 9.8 \times 10^{-10}$ for 006B).

(c) Per-cell N/C ratio against whole-cell Pearson r for cells with a valid ratio, colored by stimulation; circles denote plate 006A and triangles plate 006B. The two metrics correlate moderately (Spearman $\rho = 0.487$, $p = 2.6 \times 10^{-84}$, n = 1,400 for 006A; $\rho = 0.414$, $p = 3.5 \times 10^{-69}$, n = 1,644 for 006B) but report different aspects of the response.

Statistics: two-sided Mann–Whitney U tests of each stimulation against the matched DMSO control, with Benjamini–Hochberg FDR adjustment within plate and metric family; * $p < 0.05$, ** $p < 0.01$, *** $p < 0.001$, ns = not significant. n denotes individual cells pooled across fields of view and wells within a plate. N/C quantification is limited to cells with complete nuclear and cytosolic segmentation (valid fractions: 006A DMSO 53%, LPS 60%, TNFα 51%, IL-1β 50%; 006B DMSO 59%, LPS 60%, TNFα 55%, IL-1β 64%).

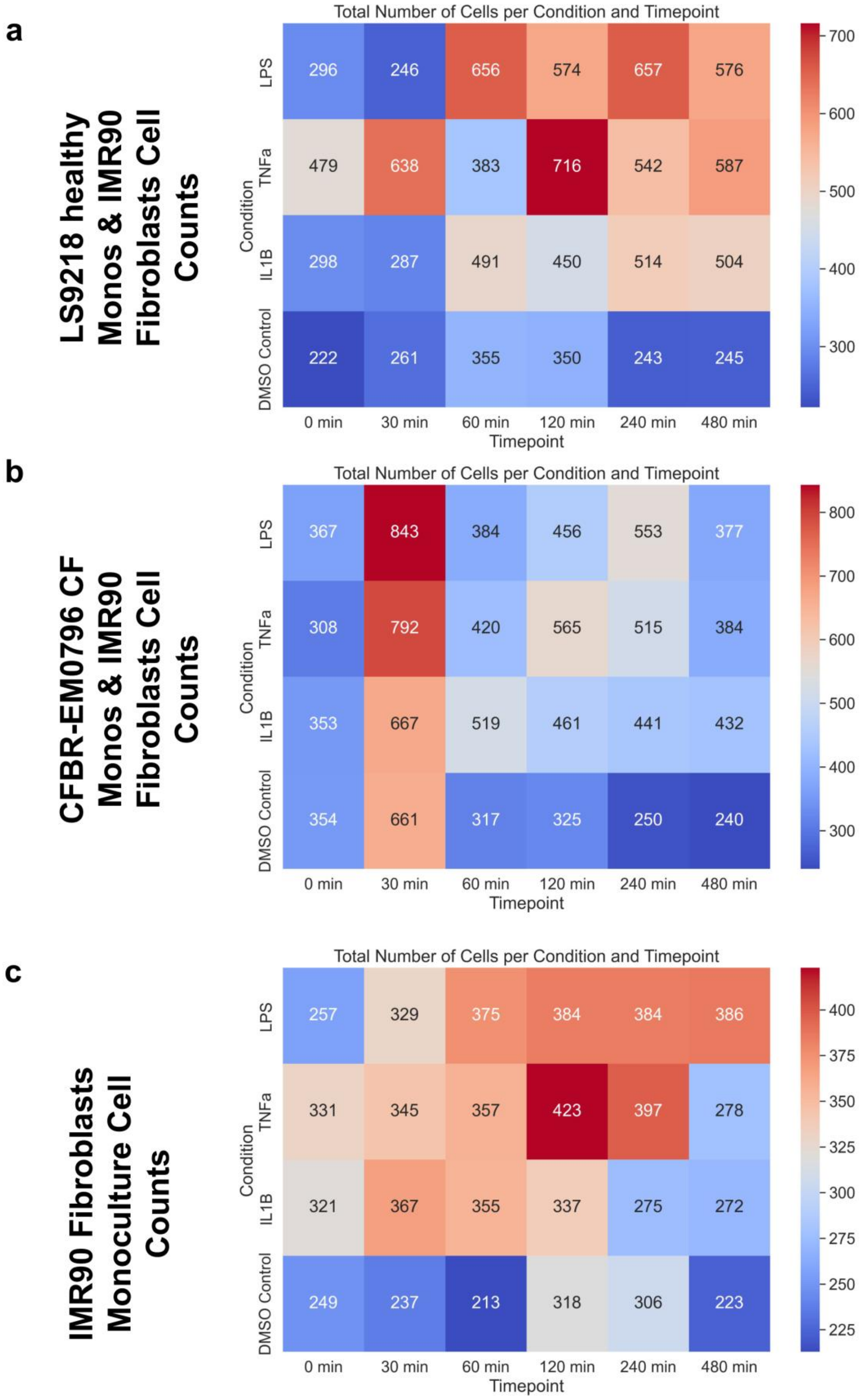
a
LS9218 healthy Monos & IMR90 Fibroblasts Cell Counts
Total Number of Cells per Condition and Timepoint
Condition
LPS
TNFa
IL1B
DMSO Control
296 246 656 574 657 576
479 638 383 716 542 587
298 287 491 450 514 504
222 261 355 350 243 245
0 min
30 min
60 min
120 min
240 min
480 min
Timepoint
700
600
500
400
300
b
CFBR-EM0796 CF Monos & IMR90 Fibroblasts Cell Counts
Total Number of Cells per Condition and Timepoint
Condition
LPS
TNFa
IL1B
DMSO Control
367 843 384 456 553 377
308 792 420 565 515 384
353 667 519 461 441 432
354 661 317 325 250 240
0 min
30 min
60 min
120 min
240 min
480 min
Timepoint
800
700
600
500
400
300
c
IMR90 Fibroblasts Monoculture Cell Counts
Total Number of Cells per Condition and Timepoint
Condition
LPS
TNFa
IL1B
DMSO Control
257 329 375 384 384 386
331 345 357 423 397 278
321 367 355 337 275 272
249 237 213 318 306 223
0 min
30 min
60 min
120 min
240 min
480 min
Timepoint
400
375
350
325
300
275
250
225

Supplementary Fig. 7. Cell count heatmaps for healthy donor monocyte co-cultures, CF patient-derived monocyte co-cultures, and corresponding IMR-90 fibroblast mono-cultures.

Heatmaps displaying the total number of segmented cells per condition and timepoint for the three iseqPLA supercomplex profiling experiments (Figs. 7–9). Each cell in the heatmap reports the number of cells identified by Cellpose nuclear segmentation within a single well. Colors are scaled from blue (low count) to red (high count) independently per panel.

**(a)** Healthy donor monocyte (donor LS9218) and IMR-90 fibroblast co-cultures (Fig. 7); n = 10,570 total cells across all conditions and timepoints.

**(b)** CF patient-derived monocyte (donor CFBR-EM0796) and IMR-90 fibroblast co-cultures (Fig. 8); n = 10,984 total cells across all conditions and timepoints.

**(c)** IMR-90 fibroblast mono-cultures without monocytes (Fig. 9); n = 7,719 total cells across all conditions and timepoints.

All three conditions were stimulated with LPS (10 ng/ml), TNFα (10 ng/ml), IL-1β (1 ng/ml), or DMSO vehicle control for 0, 30, 60, 120, 240, and 480 minutes before fixation and iseqPLA processing. These heatmaps serve as quality control metrics to confirm adequate cell sampling across all experimental conditions and to identify any condition-dependent effects on cell adhesion or detachment, paralleling the cell count heatmaps presented in Supplementary Fig. 1 for the phospho-p65 co-culture and fibroblast mono-culture experiments.

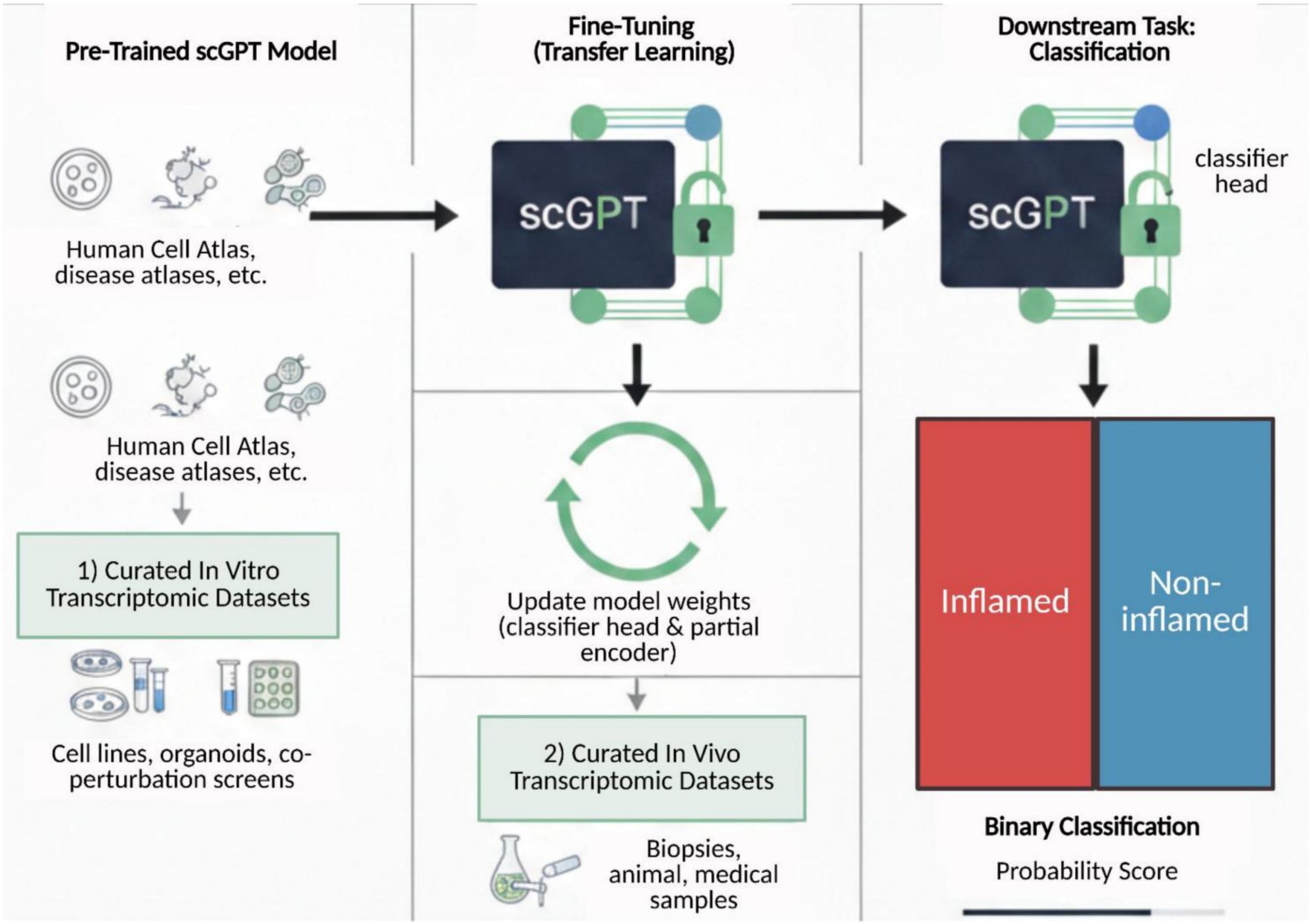
Pre-Trained scGPT Model
Fine-Tuning
(Transfer Learning)
Downstream Task:
Classification
Human Cell Atlas,
disease atlases, etc.
scGPT
scGPT
classifier
head
Human Cell Atlas,
disease atlases, etc.
1) Curated In Vitro
Transcriptomic Datasets
Cell lines, organoids, co-
perturbation screens
Update model weights
(classifier head & partial
encoder)
2) Curated In Vivo
Transcriptomic Datasets
Biopsies,
animal, medical
samples
Inflamed
Non-
inflamed
Binary Classification
Probability Score

Supplementary Fig. 8. Sequential transfer learning design of the scGPT foundation model for binary inflammatory classification.

Schematic overview of the two-stage fine-tuning pipeline applied to the pretrained whole-human scGPT transformer model. Left panel: The base scGPT model, pretrained on large-scale human cell atlas and disease atlas transcriptomic data, serves as the foundational encoder. Two curated fine-tuning corpora are prepared in parallel: (1) curated in vitro transcriptomic datasets comprising cell lines and co-perturbation screens with defined inflammatory stimuli (Supplementary Table 1), and (2) curated in vivo transcriptomic datasets spanning multiple human and mouse inflammatory lung disease cohorts (Supplementary Table 2). Center panel: Fine-tuning proceeds via transfer learning in sequential stages (first on the in vitro corpus, then on the in vivo corpus) updating both the classifier head and a partial subset of encoder weights to adapt the model to inflammation-specific transcriptional programs while preserving generalizable gene representations learned during pretraining. Right panel: The downstream classification task produces a binary output probability score distinguishing inflamed from non-inflamed cell states, enabling quantitative assessment of inflammatory gene panel relevance. Results from the in vitro and in vivo trained models are presented in Fig. 10 and Supplementary Fig. 9, respectively. Created with BioRender.com.

### In Vivo Training

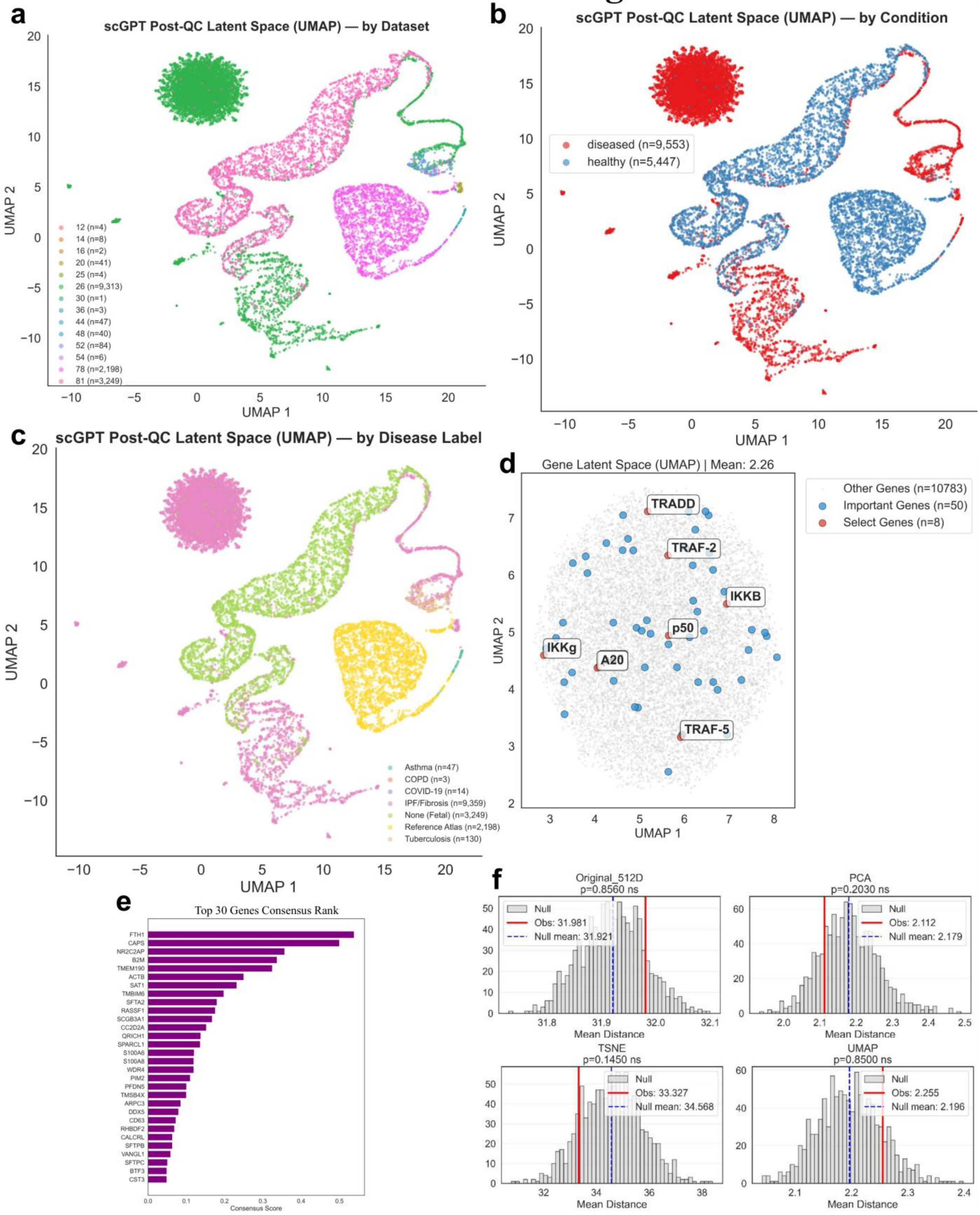

Supplementary Fig. 9. In vivo scGPT foundation model exploratory analysis of NFκB gene panel relevance across multi-disease lung datasets.

**(a–c)** UMAP visualizations of the curated in vivo transcriptomic training corpus, drawn from 15 lung-disease cohorts across human and mouse (Supplementary Table 2; n = 15,000 post-QC representative cells shown) and colored by dataset source (a), binary cell phenotype (diseased (n=9,553) vs. healthy (n=5,447)) (b), and disease label (c). The full corpus spans COVID-19 BALF, idiopathic pulmonary fibrosis (IPF), CF airway inflammation, COPD, asthma, tuberculosis, community-acquired pneumonia, and influenza, and covers scRNA-seq, CITE-seq, spatial transcriptomics, and bulk RNA-seq. Panels (a–c) show the balanced 15,000-cell subsample used for training rather than the full corpus, so only part of that range appears among the labels: the subsample is dominated by IPF/fibrosis (n=9,359), fetal reference tissue carrying no disease label (n=3,249) and reference-atlas cells (n=2,198), with tuberculosis (n=130), asthma (n=47), COVID-19 (n=14) and COPD (n=3) contributing the remainder. In (a) each dataset is identified by its numeric index in Supplementary Table 2. Starting from the in vitro fine-tuned scGPT model (Fig. 10), we further fine-tuned on this in vivo corpus with multi-GPU DataParallel training over 10 epochs, a learning rate of $5\times10^{-5}$, batch size 16 (effective per-GPU batch size 8), a maximum sequence length of 3,000 genes, and balanced class subsampling to counter the ~15:1 diseased-to-control imbalance, selecting the best model by F1 score.

**(d)** Gene latent space (UMAP) of the in vivo model, showing all genes (gray, n=10,783), the top 50 consensus-ranked reference genes (blue), and the 8 NFκB panel genes (red); the mean panel-to-reference distance was 2.26 in the UMAP space.

**(e)** Top 30 consensus gene ranking from the in vivo model, derived by aggregating importance scores across gradient-based, SHAP, and perturbation attribution methods. Highly ranked genes reflect metabolic stress and translational regulation signatures prevalent across inflammatory lung disease states.

**(f)** Permutation tests of NFκB panel-gene proximity to the top-50 reference-gene neighborhood in four distance spaces: the native 512-dimensional embedding (top left, $p = 0.8560$), the PCA projection (top right, $p = 0.2030$), the t-SNE projection (bottom left, $p = 0.1450$), and the UMAP projection (bottom right, $p = 0.8500$). In contrast to the in vitro model, none of the four tests reached significance, consistent with NFκB panel enrichment being diluted in the transcriptomically heterogeneous in vivo multi-disease corpus.

| Reference | Accession | Sequencing | Cell Type | Stimuli | Cells (pre-QC) | Cells (post-QC) | Species |
|---|---|---|---|---|---|---|---|
| Namani, A. et al. Oncotarget 8, 69847–69862 (2017) | GSE94383 | scRNA-seq (Fluidigm C1) | RAW264.7 macrophages (p65-Clover) | LPS | 823 | 818 | Mouse |
| Kull, T. et al. *Blood* 140, 99–111 (2022) | GSE199404 | scRNA-seq (TrackSeq) | HSPCs/GMPs (GFP-p65/H2B-mCherry mice) | TNFα, IL-1β | 238 | 207 | Mouse |
| Son, M. et al. *Sci Adv* 8, eabn6240 (2022) | GSE189062 | Bulk RNA-seq | 3T3 fibroblasts + macrophage co-culture | TNF, LPS (spatial gradient) | 20 | 18 | Mouse |
| Ngo, K. A. et al. *Cell Rep* 30, 2758-2775.e6 (2020) | GSE132791 | ChIP-seq (RelA), RNA-seq | MEFs (RelA KO + complementation) | TNF | 45 | 45 | Mouse |
| Derbois, C. et al. *Sci Data* 10, 433 (2023) | GSE226488 | scRNA-seq | Human PBMCs | LPS | 30,789 | 26,441 | Human |
| **Total** | | | | | 31,915 | 27,529 | |

Supplementary Table 1. Curated transcriptomic datasets for in vitro scGPT foundation model fine-tuning.

Summary of five publicly available sequencing datasets used to construct the multi-species in vitro training corpus for binary classification of control versus inflamed/treated cell states. Datasets were selected to capture NFκB-relevant cell types, stimulation conditions, and sequencing platforms in in vitro cell-line and primary-cell models, encompassing a total of 31,915 cells pre-QC (1,126 mouse + 30,789 human) and 27,529 cells after per-dataset MAD-based quality control (1,088 mouse + 26,441 human). Included datasets span RAW264.7 macrophages with LPS stimulation and a p65-Clover reporter (GSE94383, Fluidigm C1 scRNA-seq, Mouse), hematopoietic stem and progenitor cells and granulocyte-monocyte progenitors from GFP-p65/H2B-mCherry reporter mice stimulated with TNFα and IL-1β (GSE199404, TrackSeq scRNA-seq, Mouse), 3T3 fibroblast–macrophage co-cultures stimulated with TNF and LPS spatial gradients (GSE189062, bulk RNA-seq, Mouse), mouse embryonic fibroblasts with RelA knockout and complementation under TNF stimulation (GSE132791, RNA-seq with RelA ChIP-seq, Mouse), and human peripheral blood mononuclear cells stimulated with LPS (GSE226488, scRNA-seq, Human). Mouse and human vocabularies were intersected case-insensitively with an optional MGI-ortholog-based alternative, and mitochondrial (MT-prefix), cytoplasmic ribosomal (HGNC group 1054), and canonical hemoglobin (HGNC group 940) gene families were removed via HGNC-curated allowlists before training. The pretrained whole-human scGPT model was fine-tuned on this corpus to classify control versus inflamed samples; gene importance analysis and NFκB panel enrichment results are presented in Fig. 10.

| Reference | Description | Species | Modality | Accession | Num Cells |
|---|---|---|---|---|---|
| Neidleman, J. et al. Cell Reports 36, 109414 (2021) | COVID-19 BALF scRNA-seq | Human | scRNA-seq | GSM4339769 | 6249 |
| Melms, J. C. et al. Nature 595, 114–119 (2021) | COVID-19 Multi-cohort scRNA-seq | Human | scRNA-seq | GSE171524, GSE171668, GSE149878, GSE161382, GSE163919 | 10722 |
| Mayr, C. H. et al. Science Advances 10, eadl5473 (2024) | IPF Visium + Xenium Multi-modal | Human | Spatial | GSE128169, GSE128033, GSE135893, GSE122960, GSE136831, GSE158127 | 57787 |
| Reyfman, P. A. et al. Am J Respir Crit Care Med 199, 1517–1536 (2019) | Reyfman IPF Study | Human | scRNA-seq | GSE122960 | 5437 |
| Jia, M. et al. Aging Cell 22, e13969 (2023); Morse, C. et al. Eur Respir J 54, 1802441 (2019); Cruz, T. et al. J Immunol 211, 1073–1081 (2023) | Morse IPF Macrophages | Human | scRNA-seq | GSE128033 | 737280 |
| Zhang, Y. et al. Commun Biol 8, 1353 (2025) | COPD T Cell Emphysema Study | Human | scRNA-seq | GSE302339 | 4323 |
| Vieira Braga, F. A. et al. Nat Med 25, 1153–1163 (2019) | Asthma Cell Census | Human | scRNA-seq | GSE130148 | 10360 |
| Thompson, D. A. et al. Sci Rep 15, 6844 (2025) | Pediatric Obesity Asthma | Human | scRNA-seq + Bulk | GSE274583, GSE227915 | 99 |
| Alladina, J. et al. Sci Immunol 8, eabq6352 (2023); Siddiqui, S. et al. JCI Insight 6, | Asthma AEC scRNA-seq | Human | scRNA-seq | GSE164015, GSE193816 | 20410 |

| | | | | | |
|---|---|---|---|---|---|
| e139019, 139019 (2021) | | | | | |
| Wang, L. et al. J Infect 87, 373–384 (2023) | Human TB Lung scRNA-seq | Human | scRNA-seq | GSE192483 | 7507 |
| Pisu, D. et al. J Exp Med 218, e20210615 (2021); Russell, D. et al. Res Sq rs.3.rs-3934768 (2024); Pisu, D. et al. Nat Commun 15, 8522 (2024) | Mouse TB Lung CITE-seq | Mouse | scRNA-seq + CITE-seq | GSE167232 | 12 |
| Schupp, J. C. et al. Am J Respir Crit Care Med 202, 1419–1429 (2020); Öz, H. H. et al. Cell Rep 41, 111797 (2022) | CF Airway Inflammation | Human | scRNA-seq | GSE145360 | 21258 |
| Schuurman, A. R. et al. Elife 10, e69661 (2021) | Community-Acquired Pneumonia CITE-Seq | Human | CITE-Seq | GSE164948 | 9630 |
| Niethamer, T. K. et al. Cell Stem Cell 32, 302-321.e6 (2025) | Influenza ALI Longitudinal | Mouse | scRNA-seq + Spatial | GSE262927 | 3956 |
| Kasmani, M. Y. et al. Nat Commun 14, 6597 (2023) | Aging + Influenza Spatial | Mouse | scRNA-seq + Spatial + Bulk | GSE202325, GSE202322, GSE202324 | 1986 |

Supplementary Table 2. Curated transcriptomic datasets for in vivo scGPT foundation model fine-tuning.

Summary of fifteen publicly available in vivo transcriptomic datasets used to construct the multi-disease lung inflammation training corpus for binary classification of healthy versus diseased cell states. Datasets span human and mouse species across multiple inflammatory lung disease contexts, sequencing modalities (scRNA-seq, CITE-seq, spatial transcriptomics, and bulk RNA-seq), and a total of approximately 10,000 representative cells after balanced class subsampling. Included datasets cover COVID-19 bronchoalveolar lavage fluid (Neidleman et al., GSM4339769, n=6,249; Melms et al., GSE171524/171668/149878/161382/163919, n=10,722); idiopathic pulmonary fibrosis (IPF) with multi-modal Visium and Xenium spatial profiling (Mayr et al., multiple GEO accessions, n=57,787), IPF scRNA-seq (Reyfman et al., GSE122960, n=5,437), and IPF macrophage scRNA-seq (Morse et al., GSE128033, n=737,280); COPD T cell emphysema scRNA-seq (Zhang et al., GSE302339, n=4,323); asthma cell census scRNA-seq (Vieira Braga et al., GSE130148, n=10,360), pediatric obesity-associated asthma scRNA-seq and bulk RNA-seq (Thompson et al., GSE274583/227915, n=99), and asthma airway epithelial cell scRNA-seq (Alladina et al. and Siddiqui et al., GSE164015/193816, n=20,410); human TB lung scRNA-seq (Wang et al., GSE192483, n=7,507); mouse TB lung CITE-seq (Pisu et al. and Russell et al., GSE167232, n=12); CF airway inflammation scRNA-seq (Schupp et al. and Öz et al., GSE145360, n=21,258); community-acquired pneumonia CITE-seq (Schuurman et al., GSE164948, n=9,630); and influenza air-liquid interface longitudinal scRNA-seq with spatial profiling (Niethamer et al., GSE262927, n=3,956) and aging plus influenza spatial and bulk profiling (Kasmani et al., GSE202325/202322/202324, n=1,986). The in vivo fine-tuned model, built by sequential transfer learning from the in vitro model, was used to assess NF-κB panel gene enrichment within inflammation-relevant transcriptional feature space; results are presented in Supplementary Fig. 9.

| gene | consensus_score | gradient_score | gradient_score_norm | shap_score | shap_score_norm | avg_prob_change | avg_prob_change_norm |
|---|---|---|---|---|---|---|---|
| PLCG2 | 0.947254 | 3.509396 | 1 | 1.154648 | 1 | 0.18498 | 0.841763 |
| SDC2 | 0.453253 | 1.177781 | 0.335608 | 0.027887 | 0.024152 | 0.219753 | 1 |
| LMNA | 0.266965 | 0.506566 | 0.144346 | 0.047919 | 0.041501 | 0.135159 | 0.615049 |
| CCL5 | 0.195097 | 1.448484 | 0.412744 | 0.070823 | 0.061337 | 0.02444 | 0.11121 |
| SRSF5 | 0.178516 | 1.192537 | 0.339813 | 0.15487 | 0.134128 | 0.01354 | 0.061609 |
| S100A11 | 0.167257 | 0.77319 | 0.22032 | 0.096313 | 0.083413 | 0.04352 | 0.198037 |
| ANXA1 | 0.125798 | 0.800696 | 0.228158 | 0.050876 | 0.044062 | 0.023114 | 0.105175 |
| PASK | 0.115043 | 0.843382 | 0.240321 | 0.013121 | 0.011363 | 0.020536 | 0.093445 |
| ZFP36L1 | 0.090655 | 0.459758 | 0.131008 | 0.042184 | 0.036534 | 0.022948 | 0.104422 |
| CDC42 | 0.08646 | 0.790028 | 0.225118 | 0.021393 | 0.018528 | 0.003459 | 0.015734 |
| FBLN2 | 0.069613 | 0.732899 | 0.208839 | 0 | 0 | 0 | 0 |
| POLR1B | 0.066667 | 0.414164 | 0.118016 | 0.000375 | 0.000325 | 0.017946 | 0.08166 |
| CEBPA | 0.065912 | 0.566274 | 0.161359 | 0.000938 | 0.000812 | 0.007817 | 0.035564 |
| ZFP36 | 0.062995 | 0.594242 | 0.169329 | 0.011625 | 0.010068 | 0.002108 | 0.009588 |
| EOMES | 0.058772 | 0.388585 | 0.110727 | 0.034021 | 0.029464 | 0.00794 | 0.036126 |
| PTMA | 0.056147 | 0.221052 | 0.062989 | 0.076837 | 0.066546 | 0.008551 | 0.038907 |
| COL1A1 | 0.055867 | 0.588178 | 0.167601 | 0 | 0 | 0 | 0 |

| | | | | | | | |
|---|---|---|---|---|---|---|---|
| TAGLN | 0.054512 | 0.573908 | 0.163535 | 0 | 0 | 0 | 0 |
| TMSB10 | 0.052062 | 0.216809 | 0.06178 | 0.06209 | 0.053774 | 0.008931 | 0.040633 |
| DDHD1 | 0.046507 | 0.445885 | 0.127055 | 0.001044 | 0.000904 | 0.002542 | 0.011561 |
| OAZ1 | 0.046398 | 0.264984 | 0.075507 | 0.01795 | 0.015546 | 0.010581 | 0.048142 |
| IL1RN | 0.042037 | 0.301159 | 0.085815 | 0.000249 | 0.000216 | 0.008809 | 0.040081 |
| TRIOBP | 0.040889 | 0.428526 | 0.122108 | 0.000646 | 0.00056 | 0 | 0 |
| H3F3B | 0.040477 | 0.163527 | 0.046597 | 0.054506 | 0.047206 | 0.006073 | 0.027629 |
| KCNK6 | 0.039176 | 0.010842 | 0.003089 | 0.000209 | 0.000181 | 0.02511 | 0.114259 |
| AIF1 | 0.038629 | 0.146458 | 0.041733 | 0.009862 | 0.008541 | 0.01442 | 0.065613 |
| COL1A2 | 0.037913 | 0.399159 | 0.11374 | 0 | 0 | 0 | 0 |
| TNRC6C | 0.037883 | 0.349203 | 0.099505 | 0.000957 | 0.000829 | 0.002928 | 0.013316 |
| POLR3A | 0.037509 | 0.004326 | 0.001233 | 0 | 0 | 0.024458 | 0.111294 |
| NABP2 | 0.036731 | 0.008263 | 0.002354 | 0.00062 | 0.000537 | 0.023581 | 0.107302 |
| TGFBI | 0.036647 | 0.232672 | 0.0663 | 0.008196 | 0.007098 | 0.008032 | 0.036543 |
| FN1 | 0.036635 | 0.114446 | 0.032611 | 0.004138 | 0.003583 | 0.016199 | 0.073709 |
| MS4A7 | 0.036038 | 0.03761 | 0.010717 | 0.001164 | 0.001009 | 0.021183 | 0.096389 |
| CNOT11 | 0.03424 | 0.161475 | 0.046012 | 0.000149 | 0.000129 | 0.012435 | 0.056578 |
| ACTB | 0.033725 | 0.135106 | 0.038498 | 0.042488 | 0.036798 | 0.005688 | 0.025879 |
| FTH1 | 0.033597 | 0.080443 | 0.022922 | 0.04629 | 0.04009 | 0.008303 | 0.037778 |

| | | | | | | | |
|---|---|---|---|---|---|---|---|
| IFI30 | 0.033189 | 0.16472 | 0.046937 | 0.008658 | 0.007499 | 0.009919 | 0.045132 |
| PEX1 | 0.03276 | 0.002185 | 0.000623 | 0.000646 | 0.00056 | 0.021339 | 0.097098 |
| NUP93 | 0.032269 | 0.004497 | 0.001281 | 0.000115 | 1.00E-04 | 0.020971 | 0.095424 |
| IKBIP | 0.031817 | 0.005522 | 0.001573 | 0.000386 | 0.000335 | 0.020558 | 0.093543 |
| BGN | 0.031474 | 0.331363 | 0.094422 | 0 | 0 | 0 | 0 |
| BTG3 | 0.031472 | 0.173924 | 0.049559 | 0.00013 | 0.000113 | 0.009834 | 0.044745 |
| CXCL2 | 0.031461 | 0.172228 | 0.049076 | 6.75E-05 | 5.84E-05 | 0.009945 | 0.045248 |
| KLHL28 | 0.031345 | 0.00925 | 0.002636 | 0.000726 | 0.000629 | 0.019948 | 0.09077 |
| TPT1 | 0.031223 | 0.090803 | 0.025874 | 0.052609 | 0.045563 | 0.004887 | 0.022232 |
| NUDT18 | 0.030862 | 0.003968 | 0.001131 | 0.000348 | 0.000301 | 0.020032 | 0.091153 |
| KIF1C | 0.030784 | 0.006044 | 0.001722 | 0.000376 | 0.000325 | 0.019846 | 0.090305 |
| PRIM1 | 0.030093 | 0.00496 | 0.001413 | 0.001333 | 0.001154 | 0.019276 | 0.087712 |
| MYL6 | 0.030051 | 0.13683 | 0.038989 | 0.023461 | 0.020319 | 0.006779 | 0.030844 |
| EEF1A1 | 0.029971 | 0.094541 | 0.026939 | 0.05141 | 0.044524 | 0.004056 | 0.018451 |

Supplementary Table 3. Top 50 consensus-ranked discriminative genes in the fine-tuned in vitro scGPT model, used as the reference gene set for NFκB panel enrichment analysis in the scGPT gene latent space.

Ranked list of the 50 genes identified as most informative for distinguishing control from inflamed/treated cell states in the fine-tuned in vitro scGPT model (27,529 post-QC cells; datasets summarized in Supplementary Table 1). For each gene, a consensus score was computed by equally weighting three post hoc attribution methods applied to the trained classifier: (1) gradient-based attribution (gradient of the classification loss with respect to the gene embedding inputs); (2) SHapley Additive exPlanations (SHAP; Lundberg & Lee, NeurIPS 2017) estimating each gene's marginal contribution to the prediction via Shapley value approximation; and (3) perturbation analysis, which measures the change in predicted probability when individual gene inputs are masked. Each attribution method's per-gene score was min-max normalized to its corpus-wide maximum, and the consensus score was computed as the equally weighted sum of the three normalized scores such that consensus = (gradient_norm + shap_norm + perturbation_norm) / 3. Genes are ordered by descending consensus score, with values ranging from 0.9473 (PLCG2) at rank 1 to 0.0300 (EEF1A1) at rank 50. The top 30 entries of this list are displayed as the bar chart in Fig. 10d. This ranked set also constituted the "important genes" reference neighborhood against which the spatial proximity of the NFκB panel genes was evaluated in four distance spaces (the 512-dimensional scGPT gene embedding and its PCA, t-SNE, and UMAP projections) using permutation testing against random reference-set resampling (Fig. 10e,f; one reference gene lacked a valid embedding coordinate and is shown as n = 49 in panel e). Top-ranked entries span diverse inflammation-relevant functional categories including canonical NFκB-responsive chemokines and cytokines (CCL5, CXCL2, IL1RN), innate immune regulators (S100A11, ANXA1, AIF1, IFI30, MS4A7), immediate-early and AP-1/NFκB-adjacent transcriptional regulators (ZFP36, ZFP36L1, CEBPA, EOMES, BTG3), extracellular matrix and fibrotic remodeling genes (COL1A1, COL1A2, FBLN2, FN1, TGFBI, BGN, SDC2), cytoskeletal and structural genes (LMNA, TAGLN, ACTB, MYL6, TMSB10, TRIOBP), ferritin subunits (FTH1), and signaling components (PLCG2, CDC42, PASK). IKBIP (IKBKB-interacting protein, rank 40) appears in the list as a functionally NFκB-proximal regulator that is not itself part of the iseqPLA panel, providing an example of the kind of pathway-adjacent target this computational framework can nominate for future panel expansion. No mitochondrial, ribosomal, or hemoglobin genes appear in the top 50, reflecting the upstream HGNC-curated artifact-gene removal prior to training.
